# Ketogenic diet therapy for pediatric epilepsy is associated with alterations in the human gut microbiome that confer seizure resistance in mice

**DOI:** 10.1101/2023.03.17.533243

**Authors:** Gregory R. Lum, Sung Min Ha, Christine A. Olson, Montgomery Blencowe, Jorge Paramo, Beck Reyes, Joyce H. Matsumoto, Xia Yang, Elaine Y. Hsiao

## Abstract

The gut microbiome modulates seizure susceptibility and the anti-seizure effects of the ketogenic diet (KD) in animal models, but whether these relationships translate to KD therapies for human drug-resistant epilepsy is unclear. Herein, we find that the clinical KD shifts the function of the gut microbiome in children with refractory epilepsy. Colonizing mice with KD-associated human gut microbes confers increased resistance to 6-Hz psychomotor seizures, as compared to colonization with gut microbes from matched pre-treatment controls. Parallel analysis of human donor and mouse recipient metagenomic and metabolomic profiles identifies subsets of shared functional features that are seen in response to KD treatment in humans and preserved upon transfer to mice fed a standard diet. These include enriched representation of microbial genes and metabolites related to anaplerosis, fatty acid beta-oxidation, and amino acid metabolism. Mice colonized with KD-associated human gut microbes further exhibit altered hippocampal and frontal cortical transcriptomic profiles relative to colonized pre-treatment controls, including differential expression of genes related to ATP synthesis, glutathione metabolism, oxidative phosphorylation, and translation. Integrative co-occurrence network analysis of the metagenomic, metabolomic, and brain transcriptomic datasets identifies features that are shared between human and mouse networks, and select microbial functional pathways and metabolites that are candidate primary drivers of hippocampal expression signatures related to epilepsy. Together, these findings reveal key microbial functions and biological pathways that are altered by clinical KD therapies for pediatric refractory epilepsy and further linked to microbiome-induced alterations in brain gene expression and seizure protection in mice.

## INTRODUCTION

The low-carbohydrate, high-fat ketogenic diet (KD), is a mainstay treatment for refractory epilepsy, particularly in children who do not respond to existing anti-epileptic drugs. The efficacy of the KD is supported by multiple retrospective and prospective studies, which estimate that ∼30% of pediatric patients become seizure-free and ∼60% experience substantial benefit with >50% reduction in seizures (Coppola et al., 2002; Freeman et al., 1998; Hoon et al., 2005; Neal et al., 2008). However, use of the KD for treating pharmacoresistant epilepsy remains low due to difficulties with implementation, dietary compliance, and adverse side effects (Kossoff et al., 2018). Even with successful seizure reduction, retention of epileptic children on the KD is a reported 13% by the third year of dietary therapy (Hemingway et al., 2001). The primary reasons cited for discontinuation include “diet restrictiveness” and “diet side effects,” in addition to low diet responsiveness. While many pioneering studies have proposed important roles for immunosuppression, ketone bodies, anaplerosis, and gamma-aminobutyric acid (GABA) modulation in mediating the neuroprotective effects of the KD, they do not fully account for the clinical heterogeneity in patient responsiveness. Exactly how the KD confers protection against epilepsy in individuals with varied seizure semiologies remains unclear, and the biological determinants of patient responsiveness to the KD are poorly understood.

The gut microbiome plays an integral role in mediating effects of diet on multiple aspects of host physiology, including metabolism, neural activity, and behavior (Singh et al., 2017; Sonnenburg & Bäckhed, 2016). To date, a few clinical studies have reported associations between the KD regimens and alterations in the composition and/or functional potential of the gut microbiota in epileptic individuals (Lindefeldt et al., 2019; G. Xie et al., 2017; Y. Zhang et al., 2018). While promising, thus far there is little consistency across these different reports in the specific microbial taxa or gene pathways that correlate with the KD. Moreover, functional consequences of the KD- associated human epilepsy microbiome on host seizure susceptibility remain unknown. We previously reported that KD-induced alterations in the gut microbiome were necessary and sufficient for mediating the seizure protective effects of KD chow in two mouse models for refractory epilepsy – the 6-Hz psychomotor seizure model and the *Kcna1* deficiency model for sudden unexpected death in epilepsy (SUDEP) (Olson et al., 2018). Similarly, in a rat injury model of infantile spasms, transfer of the KD-induced gut microbiota into naïve animals fed a control diet reduced spasms (Mu et al., 2022). In addition, taxonomic differences in the gut microbiome were correlated with seizure severity and seizure protection in response to KD chow in the *Scn1a* deficiency model for Dravet syndrome (Miljanovic & Potschka, 2021). Together, these findings across various seizure models provide proof-of-principle that the KD alters the gut microbiome in ways that can promote seizure protection. However, whether these results from rodent studies apply to human epilepsy, the human gut microbiome, and clinical KD regimens used to treat pediatric epilepsy is still unknown, and the core microbial functions that impact seizure susceptibility are unclear.

Herein, we perform a prospective study of KD interventions in children with refractory epilepsy and test causal effects of the human gut microbiome before and after initiating clinical KD regimens on seizure susceptibility in mice. We evaluate functional changes in the human gut microbiome that are associated with KD treatment in pediatric epilepsy patients. We further identify select features of the clinical KD-associated gut microbiome that are shared across both human donors and inoculated mouse recipients that correlate with microbiome-dependent seizure protection in mice. Finally, we identify key network interactions between the gut microbiome, metabolites, and brain transcriptome that may contribute to the ability of the clinical KD-associated human gut microbiome from pediatric epilepsy patients to promote seizure protection in mice.

## RESULTS

### Clinical KD regimens elicit shared functional features of the gut microbiome in a cohort of children with refractory epilepsy

The KD is commonly prescribed for pediatric refractory epilepsy, wherein children consume commercial ketogenic infant formula and/or fat-rich, carbohydrate-restricted meals with dietary guidance from clinicians and registered dieticians (Kossoff et al., 2018). Notably, treatment regimens for the KD vary from patient to patient. KD composition depends on patient tolerability, which dictates the ratio of fat intake relative to carbohydrate and protein. Additionally, variable food sources determine the specific macro- and micro-nutrients that comprise ketogenic meals. Moreover, the KD is prescribed broadly for various forms of refractory epilepsy, the treatment population varies in genetic risk, seizure semiology, and past anti-epileptic drug exposures, among other factors. In order to assess effects of clinically-relevant KD treatments for refractory epilepsy on the gut microbiome, we therefore conducted a prospective study of 10 children with pediatric refractory epilepsy who were newly enrolled to the Ketogenic Diet Program at UCLA Mattel Children’s Hospital (**Table S1**). From each patient, we collected a stool sample within 1 day before initiating a KD regimen (pre-KD sample) and after approximately 1 month of adherence to a clinically-guided KD (post-KD sample). 1 month was chosen as a time point at which we expected to observe stabilized microbial responses to the dietary regimen (David et al., 2014).

Data from 16S rRNA gene amplicon sequencing of fecal samples indicated no significant difference in bacterial α-diversity in the post-KD fecal microbiota from pediatric epilepsy patients relative to their matched pre-KD internal controls (**Figure S1A, Table S2**). Principal coordinates analysis of unweighted and weighted Unifrac distances revealed substantial variation across individuals in baseline composition of the pre-KD microbiota (**Figure S1B**). Additionally, the clinical KD elicited differential shifts in bacterial β-diversity and varied responses across post-KD samples relative to their matching pre-KD controls, which were not significantly associated with demographic or clinical measures, such as age, sex, and prior anti-epileptic drug exposure (**Figure S1B**, Table S1, Yassour et al., 2016). Consistent with the inter-individual variation in microbial taxonomic profiles, ANCOM and ANOVA analyses (paired or unpaired) identified no significant differences in relative abundances in particular bacterial taxa when considering all post-KD sample relative to their matched pre-KD controls (**Figures S1C)**. These results indicate that, within this particular study cohort, there are no shared effects of the clinical KD on the microbial composition of the gut microbiota of children with refractory epilepsy.

Functional redundancy is common across different microbial species of the human gut microbiota (Tian et al., 2020). In light of the varied bacterial taxonomic profiles at baseline and in response to dietary treatment, we next asked whether the clinical KD is associated with shared alterations in the functional potential of the gut microbiota from children with pediatric epilepsy. Shotgun metagenomic profiling and pathway analysis indicated that compared to pre-KD samples, post-KD samples shared a significant decrease in relative abundance of genes belonging to the top 26 most abundant functional pathways, which together comprised >94% of the pathway diversity detected (**Figure S1D and S1E, Table S3**). This corresponded with a significant increase in the number of total observed pathways in post-KD samples compared to their respective pre-KD controls (**Figure S1F**). These observations suggest that the clinical KD restricts the membership of various types of microbial taxa that harbor genes related to prevalent functions and/or enriches for microbial taxa that harbor genes related to previously rare or underrepresented functions. In particular, post-KD samples exhibited significant enrichment of genes related to formaldehyde assimilation, guanosine nucleotide degradation, and L-proline biosynthesis, and decreased representation of genes related to aerobactin biosynthesis, as compared to pre-KD controls (**Figure S1G, Discussion**). There were also modest increases in genes related to GDP-mannose biosynthesis, 2-methylcitrate cycle, and glycol metabolism and degradation, and decreases in genes related to polyamine biosynthesis and biotin biosynthesis, subsets of which will be discussed in greater detail in the following sections (**Figure S1G)**. Taken together, these data suggest that treatment with KD regimens that differ in KD ratio and specific nutritional composition elicit broad shifts in the functional potential of the gut microbiome that are shared across children with varied subtypes of refractory epilepsy.

### Transferring the fecal microbiota from KD-treated pediatric epilepsy patients to mice confers seizure resistance

Causal influences of the human microbiome can be effectively studied in gnotobiotic mice, wherein transferring microbes in a clinical sample into microbiota-deficient mice is used to recapitulate the taxonomic and functional diversity of the donor human microbiota. To evaluate whether gut microbes associated with the clinical KD impact seizure susceptibility, we inoculated individual cohorts of germ-free (GF) mice with matched pre-KD and post-KD stool samples collected from children with refractory epilepsy and maintained the colonized mice on standard (non-ketogenic) mouse chow (control diet, CD). Each human donor sample (pre-KD and post-KD from 10 individuals, as biological replicates) was inoculated into 14-16 GF mice (as technical replicates) to enable cohort-level testing of susceptibility to 6-Hz psychomotor seizures (**Figure 1A**). The 6-Hz seizure model involves low-frequency corneal stimulation to induce acute complex partial seizures reminiscent of human limbic epilepsy (Barton et al., 2001). Consistent with refractory epilepsy, the 6-Hz model is resistant to several anti-epileptic drugs, but treatment with KD chow effectively protects against 6-Hz seizures in rodents (Hartman et al., 2008), raising the intensity of current required to elicit a seizure in 50% of the subjects tested (CC50, seizure threshold). The 4-day time point was chosen as the maximum duration of time that a KD-induced microbiota could be maintained in mice fed CD (Olson et al., 2018).

**Figure 1:**
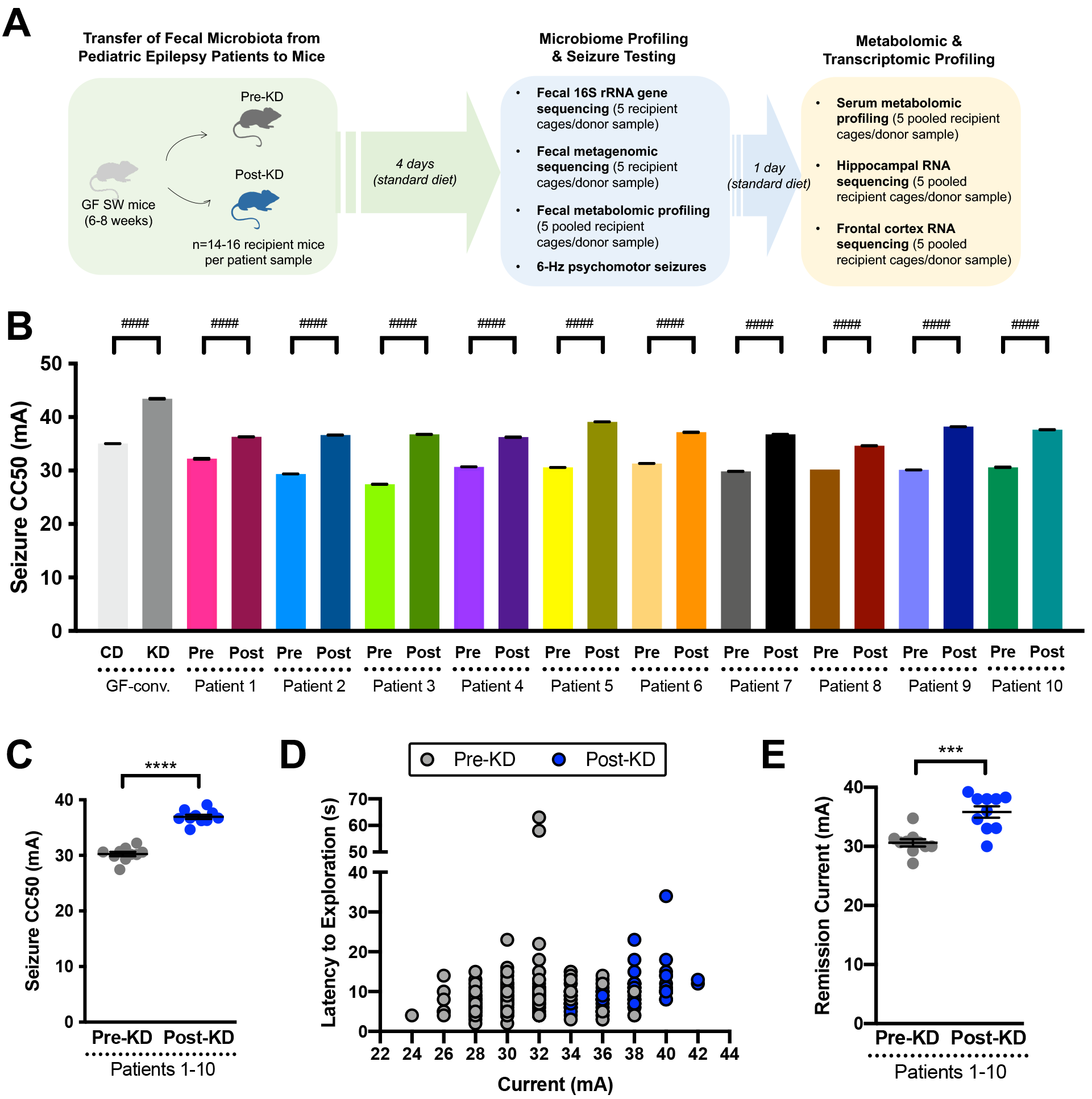
Transfer of the clinical KD-associated gut microbiota from pediatric epilepsy patients to mice confers resistance to 6-Hz seizures. **(A)** Experimental schematic for transplantation of human donor fecal microbiota samples into germ-free (GF) Swiss Webster mice for 6-Hz psychomotor seizure testing. **(B)** 6-Hz seizure thresholds for replicate mice transplanted with human microbiota from paired donor pre-KD and post-KD samples (One-way ANOVA with Tukey’s, n = 13-16 mice per patient sample, with # denoting statistical differences when considering within-patient recipient mice as technical replicates). **(C)** Average seizure thresholds of recipient mouse cohorts per patient donor sample. (Two-tailed, unpaired Welch’s t-test. n=10 patient samples per group). **(D)** Latency to exploration for all individual pre-KD (n=140) and post-KD (n=141) recipient mice. **(E)** Average current (mA) at which remission seizures were observed per patient donor sample (Two-tailed, unpaired Welch’s t-test. n = 10 patient samples per group). Data is displayed as mean ± SEM, unless otherwise noted. ^####^ p <0.0001 (within-patient mouse recipients), ***p < 0.001, ****p < 0.0001

Despite the variation in bacterial diversity across patient gut microbiota (**Figure S1**), we observed that GF mice colonized with microbes from the post-KD microbiota required greater intensity of current to induce 6-Hz seizures (**Figure 1B, Table S4**) as compared to controls colonized with microbes from the pre-KD microbiota. This effect was seen when comparing post-KD vs. pre-KD microbiota transfer for individual technical replicates per patient (**Figure 1B**), as well as when data were averaged across all patients (**Figures 1C and 1D**). In addition, compared to pre-KD controls, mice colonized with microbes from the post-KD microbiota required increased intensity of current to elicit one or more recurred seizures observed after the initial stimulus-induced seizure (**Figure 1E**), indicating that transfer of the post-KD human microbiota promotes resistance to both primary induced seizures and remission seizures in mice. On average, the post-KD samples raised seizure thresholds by 22.4% ± 6.4% relative to matched pre-KD controls (**Figures 1C and 1D**). This aligns with both our previously published data on the average effect size of KD chow on wildtype mice tested in the 6-Hz seizure assay (24.5%, Olson et al., 2018), and the observed 24.0% increase in seizure threshold seen in GF mice colonized with a conventional adult mouse microbiota (GF-conv) and fed a 6:1 KD chow, as compared to conventionalized controls fed a standard vitamin- and mineral-matched control diet (**Figure 1B**). Discrepancies in effect size across patient samples were largely driven by differences between responses for pre-KD controls (**Figure 1B**), suggesting that the comparatively low microbial diversity resulting from cross-host species transfer increases seizure susceptibility. Consistent with this, we previously observed that decreasing microbial diversity via antibiotic treatment reduced seizure threshold in the 6-Hz assay (Olson et al., 2018). Overall, these results indicate that inoculating mice with the clinical KD-associated human gut microbiota increases 6-Hz seizure threshold to levels similar to the effect sizes seen with direct consumption of the experimental 6:1 KD.

Human microbiota transplantation to mice involves oral inoculation with a human stool suspension, which is comprised of microbial biomass, as well as undigested food matter and secreted molecules from the host and microbiota. As such, effects seen in response to the transfer procedure could be due to the KD-associated gut microbiota or microbiota-independent dietary or host factors. To gain insight into whether bacteria from the gut microbiota are required for mediating the increases in seizure protection seen with inoculation of the human post-KD microbiota into mice, mice inoculated with a randomly selected post-KD donor sample were treated with broad-spectrum antibiotics (**Abx**) to deplete the microbiota, or with vehicle (**Veh**) as negative control (**Figures S2A and S2B**). Mice that were inoculated with the post-KD sample and treated with Veh displayed seizure thresholds that were comparable to that seen previously in recipient mice without the added Veh treatment (**Figures S2C, S2D, and 1B and Table S4**). This suggests that the post-KD sample induced increases in seizure resistance that were maintained for at least 12 days in mice fed CD. In contrast, depletion of gut bacteria in mice that were colonized with the post-KD microbiota decreased seizure thresholds to levels that were lower than previously seen in pre-KD colonized controls (**Figures S2C, S2D, and 1B**). These results indicate that bacterial members of the post-KD microbiota are necessary for mediating the increases in seizure threshold seen in response to transfer of the clinical-KD associated microbiota from a pediatric epilepsy patient into mice.

Administration of microbial metabolites or other microbiome-dependent molecules, in lieu of viable microbiota, has been reported to ameliorate symptoms of recurrent *Clostridiodes difficile* infection, inflammatory bowel disease, and multiple sclerosis, among other conditions (Cekanaviciute et al., 2017; Levy et al., 2015; Ott et al., 2017). To gain insight into whether administration of clinical KD-associated intestinal small molecules is sufficient to confer seizure protection in mice, a post-KD donor sample selected at random was sterile filtered and then administered to a cohort of GF recipient mice (**Figure S3A**), alongside controls that were administered the unfiltered post-KD suspension, as was done previously for human microbiota inoculation (**Figures S3A and 1B**). At 4 days post inoculation, mice that were treated with the post-KD filtrate exhibited lower seizure threshold compared to controls that were treated with the corresponding unfiltered post-KD suspension (**Figures S3B and S3C and Table S4**). These data indicate that clinical KD-associated small molecules in the post-KD fecal sample from a pediatric epilepsy patient are not sufficient to confer persistent seizure protection in mice.

Orally administered microbial metabolites can be rapidly absorbed and cleared from systemic circulation within a few hours of administration (Abrams & Bishop, 1967; Williams et al., 2020). To further assess whether clinical KD-associated intestinal small molecules, including microbial metabolites, acutely modulate seizure susceptibility, mice were orally gavaged with a sterile-filtered post-KD sample and assessed 2 hours later for 6-Hz seizure threshold, rather than 4 days later as in the previous experiments (**Figure S4A**). Mice treated with post-KD filtrate exhibited significantly increased seizure protection compared to controls treated with pre-KD filtrate (**Figures S4B and S4C and Table S4**), with seizure thresholds that approached those seen after inoculation of the post-KD suspension (**Figures S4B and 1B**). These data indicate that administration of clinical KD-associated intestinal small molecules can acutely confer seizure protection in mice over short timescales (i.e., 2 hours, **Figure S4**), which diminishes by 4 days post treatment (**Figure S3**). Taken together, the results presented in these series of experiments suggest that the clinical KD for pediatric refractory epilepsy is associated with alterations in metabolic activities of the gut microbiota that promote seizure resistance in mice.

While the “humanization” of mice with microbiota from clinical stool samples is a powerful tool for translational microbiome research (Turnbaugh et al., 2009), the approach has technical and biological limitations that warrant careful consideration (Walter et al., 2020). Namely, while much of the taxonomic and functional diversity of the donor inoculum can be recapitulated in recipient mice (Bokoliya et al., 2021), developmental influences and host-specific selection (Rawls et al., 2006), among other factors, preclude full “engraftment” of the human gut microbiota in GF mice (Walter et al., 2020). To evaluate the fidelity of fecal microbiota “transplantation” from pediatric epilepsy patients to GF mice, we subjected both the donor pre-KD and post-KD stool samples and corresponding recipient mouse fecal pellets collected at 4 days post-inoculation (the day of seizure testing) to 16S rRNA gene amplicon sequencing (**Figure 1A, Tables S2 and S5**). Principal coordinates analysis of bacterial taxonomic data revealed overt clustering of donor samples with matched recipient samples only for select patients, while the remaining exhibited substantial variation and no noticeable clustering (**Figure S5A**). There was no significant difference in α-diversity between pre-KD and post-KD fecal microbiota for either donor or recipient samples (**Figures S5B**). However, we observed a significant reduction in α-diversity, with an average decrease of 38% for all mouse recipient microbiota relative to all human donor microbiota (**Figure S5C**), indicating incomplete transfer or engraftment of the human microbiota in mice. These results align with several previous reports of reduced bacterial α-diversity in mice inoculated with human microbiota, with estimated decreases of 35%, 38%, and 50% (Blanton et al., 2016; Sharon et al., 2019; Staley et al., 2017), suggesting that we achieved levels of transfer fidelity that are consistent with those in the field. However, the inability to fully recapitulate the taxonomic diversity of the human gut microbiota from pediatric epilepsy patients in mice draws into question whether the increases in seizure resistance seen in mice inoculated with post-KD microbiota are relevant to the actual clinical condition. We therefore focused subsequent experiments on identifying and evaluating the subset of functional features of the KD-associated human gut microbiome that are recapitulated in recipient mice, and the microbiome-dependent alterations in host physiology that correspond with seizure protection in mice.

### Select functional features of the clinical KD-associated human microbiome are recapitulated in colonized recipient mice and correlate with seizure protection

Given the widespread use of the clinical KD for treating epilepsy, and an increasing number of other neurodevelopmental and neurodegenerative disorders, elucidating how the activity of the gut microbiome is altered by the clinical KD could reveal important insights into its physiological effects. To identify microbiome associations with the clinical KD and further determine which of the associations, if any, may modify seizure risk, we functionally characterized the gut microbiome from pediatric epilepsy patients before and after treatment with the clinical KD, as well as from gnotobiotic mice that were inoculated with the patient samples, and tested for causal outcomes on seizure susceptibility. Metagenomic sequencing and analysis revealed microbial gene pathways that were differentially abundant in post-KD samples relative pre-KD controls, and shared between both human donor samples and mouse recipient samples (**Figure 2A, Tables S3 and S6**). In particular, microbial genes relevant to fatty acid β-oxidation, glycol metabolism and degradation, methylcitrate cycle I, methylcitrate cycle II, and proline biosynthesis were similarly elevated in post-KD human samples and post-KD-inoculated mice compared to their respective pre-KD controls (**Figures 2A and 2B**). These findings align with reported influences of the KD on fatty acid oxidation (A. R. Kennedy et al., 2007), of carbohydrate restriction on promoting the glyoxylate cycle (Puckett et al., 2017), and of fatty acid β-oxidation on the initiation of the methylcitrate cycle (Clark & Cronan, 2005). Proline metabolism involves reactions with glutamine, glutamate, ornithine, and arginine, which might relate to reported effects of KD on amino acid metabolism, particularly of glutamine and glutamate (Yudkoff et al., 2007). In addition, both post-KD human donor and mouse recipient samples exhibited reductions in microbial genes relevant to polyamine biosynthesis and aerobactin biosynthesis (**Figures 2A and 2B**). The main role of polyamine biosynthesis is generation of putrescine, mainly using the glucogenic amino acid L-arginine which is consumer in reduced amounts while on the KD. Aerobactin, a sidophore, biosynthesis uses the ketogenic amino acid L-lysine, which is also essential for acetyl-CoA synthesis and energy production during ketosis. These data suggest that the consumption of a clinical KD by children with refractory epilepsy enriches for gut microbes that have the functional capacity to metabolize dietary fats and to perform anaplerotic reactions when dietary carbohydrates are restricted. The findings further indicate that these general features of the KD-associated human gut microbiome are phenocopied in recipient mice that exhibit microbiome-dependent protection against 6-Hz seizures.

**Figure 2:**
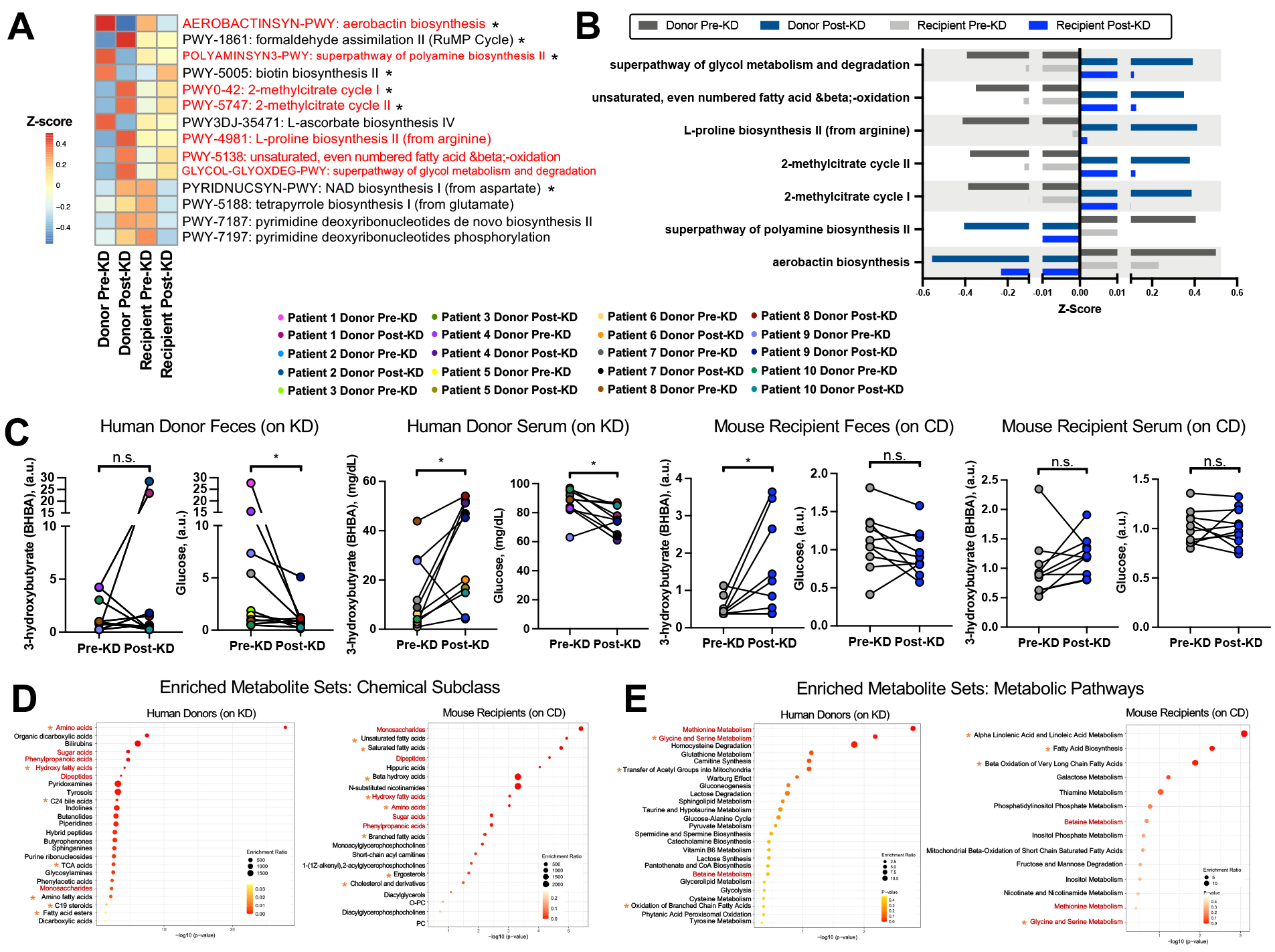
The clinical KD-associated human microbiome exhibits functional alterations that are phenocopied in seizure-protected recipient mice. **(A)** Microbial functional pathways differentially abundant as determined by MaAsLin2 analysis comparing post-KD (samples relative to pre-KD controls for either donor fecal samples or recipient fecal samples (donors: n=10 per diet condition; recipients: n=10 per donor diet condition [50 total, where each n reflects average of 5 technical replicate recipient mice per donor patient sample]). Red font denotes pathways that are commonly differentially abundant in the same direction in both donors fed KD and recipient mice fed CD (all pathways listed had minimum p<0.1; *denotes pathways with p<0.05). **(B)** Microbial functional pathways that are commonly differentially abundant by MaAsLin2 analysis in the same direction in post-KD donor and recipient controls (donors: n=10 per diet condition; recipients: n=10 per donor diet condition [50 total, where each n reflects average of 5 technical replicate recipient mice per donor patient sample]). **(C)** Beta-hydroxybutyrate (BHBA) and glucose levels in human donor (left) and mouse recipient (right) pre-KD and post-KD feces and serum (two-tailed Wilcoxon matched-pairs signed rank test; donors: n=10 per diet condition; recipients: n=10 per donor diet condition [50 total, where each n reflects average of 5 technical replicate recipient mice per donor patient sample]). **(D)** Metabolite set enrichment analysis showing the top 25 enriched chemical subclasses ordered by p-value for the set of differentially abundant metabolites in human donor (left) post-KD vs pre-KD fecal samples (p<0.05, two-tailed, matched pairs Student’s t-test, n=10 per diet condition). Metabolite set enrichment analysis showing enriched chemical subclasses ordered by p-value for the set of differentially abundant metabolites in recipient mouse (right) post-KD vs pre-KD fecal samples (p<0.05, matched pairs Student’s t-test, n=10 per patient diet condition, where each sample is pooled from 5 recipient mice per donor patient sample). Red font denotes chemical subclasses altered in post-KD vs pre-KD human donor feces that are shared with those differentially regulated in post-KD vs pre-KD mouse recipient feces. Orange asterisks (*) denote additional chemical subclasses that are relevant to KD based on existing literature. **(E)** Metabolite set enrichment analysis showing the top 25 enriched SMPBD pathways by p-value for the set of differentially abundant metabolites in human donor (left) post-KD vs pre-KD fecal samples (p<0.05, matched pairs Student’s t-test, n=10 per patient diet condition). Metabolite set enrichment analysis showing enriched SMPBD pathways by p-value for the set of differentially abundant metabolites in recipient mouse (right) post-KD vs pre-KD fecal samples (p<0.05, matched pairs Student’s t-test; n=10 per patient diet condition, where each sample is pooled from 5 recipient mice per donor patient sample). Red font denotes metabolic pathways altered in post-KD vs pre-KD human donor feces that are shared with those differentially regulated in post-KD vs pre-KD mouse recipient feces. Orange asterisks (*) denote additional chemical subclasses that are relevant to KD based on existing literature. Data is displayed as mean ± SEM, unless otherwise noted. *p < 0.05. n.s.=not statistically significant. KD, ketogenic diet; BHBA, beta-hydroxybutyrate; CD, control diet; SMPDB, The Small Molecule Pathway Database; PC=phosphatidylcholine

The observed metagenomic signatures reveal clinical KD-associated changes in the functional potential of the gut microbiome that are preserved upon transfer to GF mice. To identify clinical KD-induced alterations in the functional activity of the gut microbiome, we performed untargeted metabolomic profiling of aliquots of the same donor fecal samples from pediatric epilepsy patients collected before and after initiating the KD regimen, and of both fecal and serum samples from recipient mice that were inoculated with the pre-KD or post-KD human fecal microbiota and fed CD (**Tables S7, S8, and S9**). Results from clinical laboratory testing of human blood samples confirmed that the month-long clinical KD regimen elevated serum β-hydroxybutyrate (**BHBA**) levels and reduced serum glucose levels, relative to pre-KD concentrations, in pediatric refractory epilepsy patients (**Figure 2C**). Decreases in glucose, but not BHBA, were similarly seen in human post-KD stool samples relative to matched pre-KD controls (**Figure 2C**), which is consistent with dietary carbohydrate restriction and KD-induced BHBA synthesis by the liver to elevate systemic, but not fecal, BHBA levels (Westman et al., 2007). Transfer of the post-KD human microbiota into mice yielded no significant differences in serum BHBA or glucose relative to pre-KD recipient controls (**Figure 2C**), indicating that the clinical KD-associated microbiota does not sufficiently promote key systemic features of ketosis in mice fed the standard CD. Notably, however, mice that were inoculated with post-KD human microbiota and fed CD exhibited statistically significant increases in fecal BHBA levels relative to matched pre-KD recipient controls (**Figure 2C**). This could reflect alterations in intestinal synthesis of BHBA (Mierziak et al., 2021) and/or in microbial utilization of host-derived BHBA (Ang et al., 2020). Since this effect was not seen in the donor human fecal samples, we reasoned that this phenotype is likely an artifact of the “transplantation” approach and/or experimental design, and therefore not relevant to the clinical condition. These results suggest that transfer of the clinical KD-associated human gut microbiota into mice promotes resistance to 6-Hz seizures (**Figure 1**) through mechanisms that act independently of ketosis.

We further assessed results from untargeted metabolomic profiling to identify metabolites that were differentially abundant in post-KD samples relative to pre-KD controls and patterns that were shared across human donor and mouse recipient samples (**Figure 2D, Figure S6, Tables S7 and S8**). Despite heterogeneity in the patient population and specific clinical KD regimens, 79 metabolites were significantly differentially abundant in fecal samples from post-KD human fecal samples relative to their matched pre-KD controls (**Figure S6A, Table S7**). 336 metabolites were identified in both the human fecal samples and mice fed the 6:1 KD chow vs. vitamin- and mineral-matched control chow for 2 weeks samples, previously published by our group (**Table S10**, Olson et al., 2018). 35 metabolites were differentially regulated in human fecal samples and 169 metabolites were differentially regulated in mouse fecal samples (**Figure S6B**). Of these significantly altered metabolites, 20 were found to be changed in the same direction across human and mouse samples (**Figure S6B-D**). These included KD-induced increases in levels of metabolites related to fatty acid beta-oxidation, such as palmitoleoylcarnitine (C16:1) and oleoylcarnitine (C18:1), and a decrease in kynurenine which have previously been associated with seizure susceptibility (Żarnowska et al., 2019). This statistically significant overlap suggests that there are biochemical changes that are shared across clinical KD treatments for pediatric epilepsy and mouse models of KD, and that some of the fecal metabolomic alterations observed in KD-treated epilepsy patients are a direct consequence (rather than correlate) of dietary intervention. Of the 20 significantly differentially abundant metabolites shared in human and mouse, 14 (∼70%) were further significantly altered by antibiotic treatment to deplete gut bacteria in KD-fed mice (**Figure S6C,** Olson et al., 2018). Taken together, these data indicate that clinical KD regimens alter fecal metabolites in children with refractory epilepsy, a subset of which have the potential to be microbiome-dependent.

Although there was substantial variability in taxonomic composition of microbiota within mouse recipients of the same experimental condition (**Figure S1-S2**), fecal samples from mouse recipient cohorts exhibited statistically significant alterations in 45 metabolites that were shared when considering all post-KD mouse recipient fecal samples relative to their pre-KD controls (**Table S8, Figure S6A**). Notably, however, none of these 45 differentially abundant metabolites in mouse feces were identical to the 79 differentially abundant metabolites seen in human donor fecal samples (**Figure S6A**), which could reflect host specific metabolite utilization and the fact that recipient mice were fed standard chow (CD), while human donors were consuming a clinical KD at the time of sample collection. To gain insight into whether the differentially abundant metabolites relate to similar biological functions, we performed metabolite set enrichment analysis (MSEA) of the significantly altered metabolites in human vs. mouse (Pang et al., 2021). MSEA of the significantly altered metabolites identified select chemical classes (**Figure 2D**) and metabolic pathways (**Figure 2E**) that were similarly enriched in both human donor and mouse recipient post-KD samples relative to pre-KD controls. In particular, there was shared enrichment of amino acid, hydroxy fatty acid, sugar acid, phenylpropanoic acid, and monosaccharide-related metabolites across post-KD conditions for both human donors and mouse recipients (**Figure 2D**). Differentially abundant metabolites from human post-KD fecal samples also exhibited enrichment of bile acids and other fatty acid derivatives (**Figure 2D, left**), which might reflect KD- and/or microbiome-driven alterations in lipid metabolism (Joyce et al., 2014).

For metabolic pathways, post-KD samples for both human donor and mouse recipient conditions exhibited differential abundance of metabolites related to methionine metabolism, glycine and serine metabolism, and betaine metabolism (**Figure 2E**). While the biological relevance to KD is unclear, one possibility is that these pathways reflect known influences of the KD on one-carbon (1C) metabolism, a series of interlinking metabolic pathways that control levels of methionine, serine, and glycine, and that integrate nutrient availability with cellular nutritional status (Ducker & Rabinowitz, 2017). In addition, differentially abundant fecal metabolites from mouse post-KD recipients mapped to pathways related to alpha linolenic acid and linoleic acid metabolism, fatty acid biosynthesis, and beta-oxidation of very long chain fatty acids (**Figure 2E, right**), which aligns with the observed metagenomic enrichment in microbial genes related to fatty acid metabolism in response to the clinical KD (**Figure 2A-B**). Some of the differential metabolite chemical subclasses and metabolic pathways in mouse fecal samples were similarly seen in matched mouse serum samples (**Table S9**) – in particular, metabolites representing amino acid, hydroxy fatty acid, and unsaturated fatty acid subclasses, and related to alpha linolenic acid and linoleic acid metabolism, betaine metabolism, and beta-oxidation of fatty acids were altered in both feces and serum of mice receiving post-KD samples relative to pre-KD controls (**Figure S6E**). Taken together, these results suggest that the clinical KD induces alterations in the function of the gut microbiome of pediatric epilepsy patients, and that a subset of these functional characteristics may be phenocopied upon microbial transfer to mice, which develop microbiome-dependent resistance to 6-Hz seizures.

### Transferring the fecal microbiota from KD-treated pediatric epilepsy patients to mice induces alterations in brain gene expression

Seizures result from atypical neural function related to discharge of electrical signals or failure to constrain the spread of these signals. To gain insight into how colonization with microbes derived from the fecal microbiota of KD-treated individuals may alter brain function to modify seizure susceptibility, we performed transcriptomic profiling of brain tissues from cohorts of mice colonized with microbes from the post-KD human microbiota or pre-KD controls. We focused on the hippocampus and frontal cortex based on their relevance to human epilepsy, their involvement in initiating psychomotor seizures in the 6-Hz seizure assay, and evidence that the microbiome can alter gene expression and metabolites in these brain regions (Chauhan et al., 2022; Suarez et al., 2018). RNA sequencing of hippocampal tissues revealed many differentially expressed genes that were seen in post-KD samples relative to pre-KD controls (**Table S11**), including those related to core cell biological processes relating to RNA processing, translation, cellular stress response, TORC1 signaling, regulation of long-term synaptic potentiation, neuronal development, and response to nutrient levels (**Figure 3A**). The most drastic alterations included upregulation of *Dusp12, Bmpr1b,* and *Cmya5* and downregulation of *Abcc9*, *Ufsp1*, and *Tbx2* transcripts (**Figure 3B**). *Dusp12* is a dual specificity phosphatase that can dephosphorylate phosphothreonine and phosphoserine (Muda et al., 1999)*, Bmpr1b,* a member of the bone morphogenic receptor family, is a serine/threonine kinase influencing neuronal cell fate (Venugopal et al., 2012), and *Cmya5* encodes for myospyrn which is essential for structural integrity during neuritogenesis (Hsiung et al., 2019). *Abcc9* is an ATP-binding cassette transporter encoding the sulfonylurea receptor 2 subunit for potassium channels (Nelson et al., 2015), *Ufsp1* is a Ufm1 specific protease that regulates ubiquitin-like conjugation and has been linked to seizures (Millrine et al., 2022), and *Tbx2* is a transcription factor linked to neuronal cell cycle control and neuroinflammation (Reinhardt et al., 2019). STRING network analysis additionally revealed top protein interaction clusters enriched for essential biological processes including RNA processing, oxidative phosphorylation, and cell cycle regulation, consistent with results from GO enrichment analysis (**Figure 3A, 3C, 3D**), as well as endocytosis and glutathione metabolism (**Figure 3D**).

**Figure 3:**
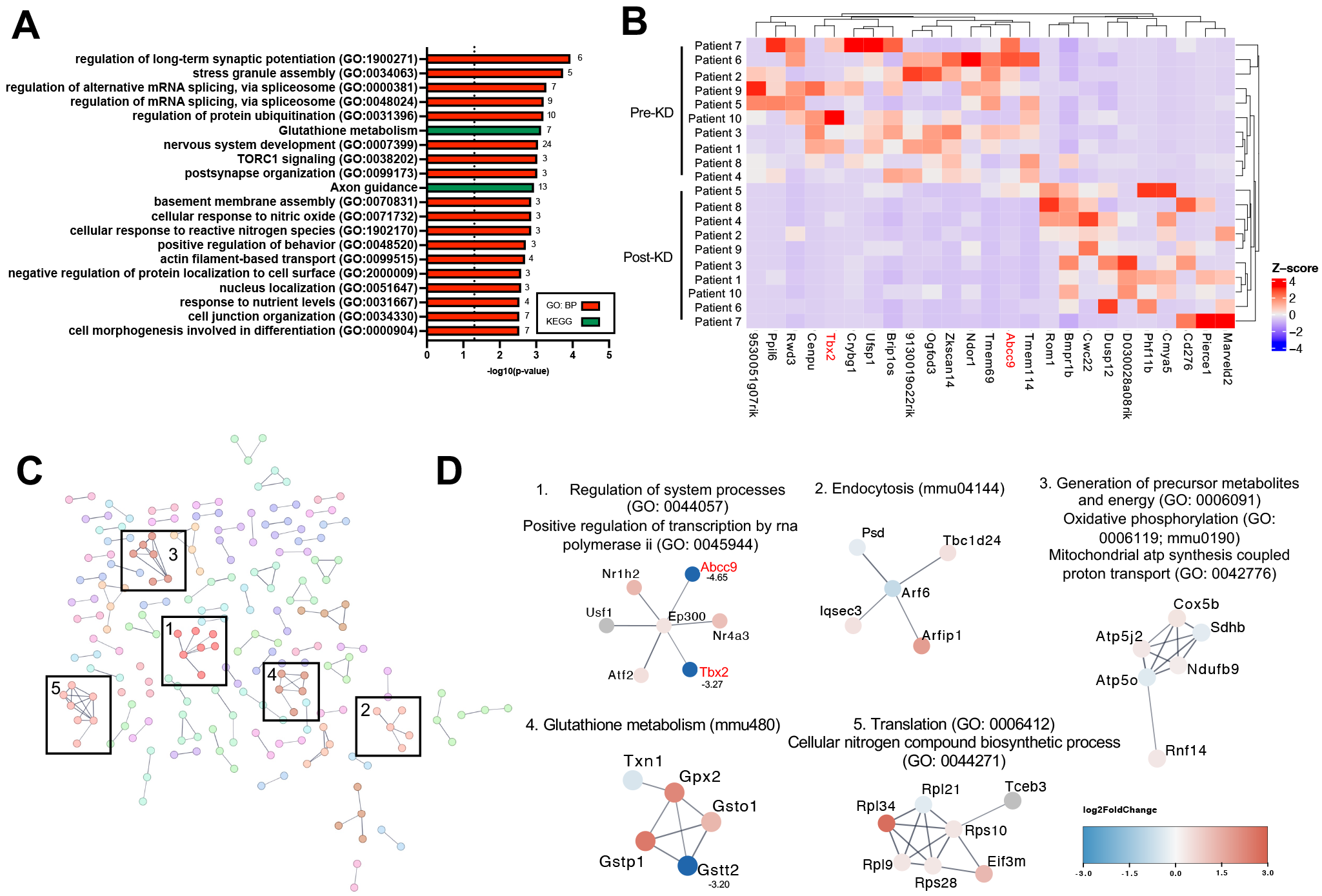
Seizure resistance in mice inoculated with the post-KD microbiota is associated with alterations in the brain transcriptome. **(A)** GO: Biological Process gene ontology of differentially expressed genes (p<0.05) in recipient mouse post-KD (n=10, where each sample is pooled from 6 recipient mice per donor patient sample) compared to pre-KD hippocampal samples, top 20 ranked by p-value (n=10 per patient diet condition, where each sample is pooled from 6 recipient mice per donor patient sample). **(B)** Heatmap of euclidian row and column clustered top 25 differentially expressed genes in recipient mouse post-KD compared to pre-KD hippocampus ranked by p-value, smallest to largest, and with log2 fold-change >2 (n=10 per patient diet condition, where each sample is pooled from 6 recipient mice per donor patient sample). **(C)** Protein interaction network with MCL clustering based upon mouse recipient post-KD and pre-KD hippocampal transcriptomics which appeared in both GO and STRING network enrichment analyses, STRING network enrichment score >0.7 (n=10 per patient diet condition, where each sample is pooled from 6 recipient mice per donor patient sample). **(D)** Functional enrichment of top MCL sub-network clusters from hippocampal transcriptomics STRING network analysis, proteins are colored based on their overall log2FC. If log2FC >3 or <-3, the value is listed next to the node name (n=10 per patient diet condition, where each sample is pooled from 6 recipient mice per donor patient sample). KD, ketogenic diet; GO, gene ontology; MCL, Markov Cluster Algorithm.

Some differentially expressed genes were also identified in frontal cortical tissues of post-KD recipients relative to pre-KD controls (**Table S12**), which similarly to hippocampus, included those related to core cell biological processes for RNA surveillance and catabolism, cellular stress responses, TORC1 signaling, and further included genes related to potassium ion transport, and core carbohydrate metabolism (**Figure S7A**). The most drastic alterations included upregulation of *Serpinb1a, Nqo1,* and *Slc6a12* transcripts, and downregulation of *Aldh3b1, Setmar*, and *Tfb1m* transcripts (**Figure S7B**). *Serpinb1a* is a serine/cysteine protease inhibitor (Huasong et al., 2015)*, Nqo1* encodes an antioxidant enzyme that primarily catalyzes the reduction of quinones (Ross & Siegel, 2021), and *Slc6a12* encodes for a betaine-GABA transporter (Zhou et al., 2012)*. Aldh3b1* is an aldehyde dehydrogenase linked to oxidative stress reduction (Marchitti et al., 2007)*, Setmar* encodes a histone-lysine N-methyltransferase (Cordaux et al., 2006), and *Tfb1m* has been shown to function as methyltransferase (Metodiev et al., 2009). STRING network clustering analysis additionally revealed top protein interaction clusters enriched for transcription regulation, translation, and oxidative phosphorylation, also seen in frontal cortex GO enrichment analysis and in the hippocampal STRING network, as well as clusters enriched for calcium signaling, transcriptional regulation, and translation (**Figure S7C, S7D**). Differentially expressed gene sets from both hippocampus and frontal cortex were enriched for TORC1 signaling, cellular response to stress, and oxidate phosphorylation through GO enrichment and STRING clustering, which have all been shown to affect seizure susceptibility (Chan, 2001; Nguyen & Bordey, 2021). The similarities between transcriptomic results from hippocampus and frontal cortex suggest that colonization with post-KD microbes elicits key alterations in host metabolism that impact core biological processes that are generally consistent across different brain regions. Overall, these results indicate that mice that acquire seizure resistance in response to colonization with microbes from the post-KD human gut microbiota exhibit alterations in hippocampal and frontal cortical gene expression, relative to pre-KD recipient controls.

### Multi’omics analysis reveals network connections linking microbial genomic pathways and metabolites to hippocampal transcripts related to epilepsy

To further identify key gut microbial functions that may drive particular brain gene expression signatures, we utilized microbe-metabolite vectors (MMVEC) (Morton et al., 2019) to build an integrated network of fecal metagenomic, fecal metabolomic, serum metabolomic, hippocampal transcriptomic, and frontal cortical transcriptomic datasets from mice inoculated with the human pre-KD or post-KD microbiota (**Table S13**). We generated a parallel network comprised of fecal metagenomic and fecal metabolomic datasets from human pre-KD and post-KD donor stool samples to identify features similarly underscored in both human and mouse networks, suggesting their clinical relevance. The human donor and mouse recipient networks were linked by 3 common nodes - metagenomic pathways describing branched chain amino acid (BCAA) biosynthesis (BRANCHED-CHAIN-AA-SYN-PWY), L-alanine fermentation (PROPFERM-PWY), and co-enzyme A biosynthesis (COA-PWY), as well as pathways for arginine synthesis (ARGSYN-PWY In the human network and ARGSYNSUB-PWY in the mouse network) (**Figure 4A**, center gray and green nodes). The shared BCAA biosynthesis, co-enzyme A biosynthesis, and arginine synthesis pathways were also identified by weighted key driver analysis as highly interconnected across the ‘omics datasets and essential regulator nodes of the network (Ding et al., 2021) (**Figure 4A**, diamonds). The human donor network also contained an additional key driver metagenomic node for isoleucine biosynthesis (ILEUSYN-PWY), which aligns with the metagenomic node for BCAA biosynthesis. Consistent with the shared metagenomic key drivers between mouse and human networks, the human fecal metabolomic module was enriched for nodes related to valine, leucine, and isoleucine (BCAA) and CoA biosynthesis (**Figure 4A**, gray diamond node). In the mouse network, metabolomic modules included nodes related to glycerophospholipid metabolism for fecal metabolites and pentose and glucuronate interconversions for serum metabolites (**Figure 4A**, orange and sea green nodes). Nodes for fecal 1-(1-enyl-palmitoyl)-2-linoleoyl-GPC (P-16:0/18:2)*, 1-(1-enyl-palmitoyl)-2-palmitoyl-GPC (P- 16:0/16:0)*, 1-(1-enyl-palmitoyl)-2-arachidonoyl-GPC (P-16:0/20:4)*, 3-hydroxybutyrate (BHBA), and myo-inositol (**Figure 4A**, orange metabolite nodes in red font) were similarly identified as differentially abundant in individual metabolomic analyses for recipient post-KD fecal samples relative to pre-KD controls (**Table S7**). These mouse metagenomic and metabolomic modules were linked to 5 transcriptomic modules for hippocampal genes and 3 for frontal cortical genes (**Figure 4A**, bottom section). The hippocampal transcript modules were enriched for nodes related to regulation of telomerase RNA localization to Cajal body, glycosylphosphatidylinositol (GPI) anchor biosynthetic processes, Wnt signaling, and neuron generation and migration (**Figure 4A**). The frontal cortical transcript modules were enriched for nodes related to regulation of catabolic processes, lipase activity, and BCAA transmembrane transporter activity (**Figure 4A**). This suggests that these particular biological processes are most closely associated with the microbial functional features identified in the network. The transcript nodes included 41 hippocampal genes and 4 frontal cortical genes that were similarly identified in individual transcriptomic analyses as differentially expressed in post-KD recipient mice relative to pre-KD controls (**Figure 4A**, transcript nodes in red font). The higher number of connections between metabolomic modules and hippocampal transcripts suggests that the gut microbiome may exhibit a greater regulatory role for the hippocampus than for the frontal cortex in post-KD recipient mice compared to pre-KD controls. Of particular interest are the links between fecal metabolites related to glycerophospholipid metabolism, which are regulated by the microbiome (Zheng et al., 2021), and hippocampal transcript modules enriched for Wnt signaling and GPI anchor biosynthetic processes, pathways implicated in seizure susceptibility (Hodges & Lugo, 2018; Wu et al., 2020).

**Figure 4:**
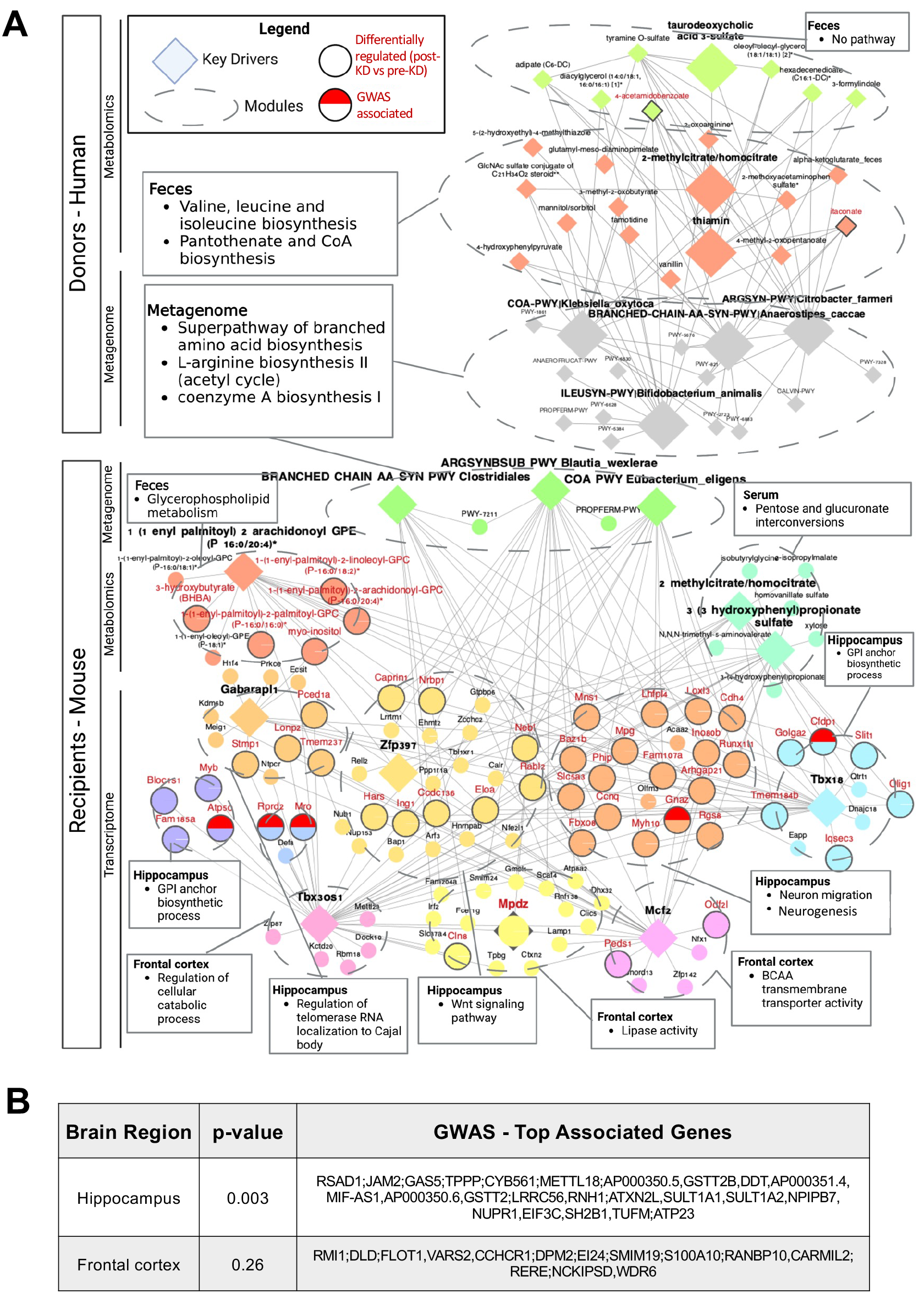
Multi’omic network analysis identifies key microbial genomic pathways and microbially modulated metabolites associated with differential expression of hippocampal transcripts. **(A)** MMVEC based co-occurrence network constructed from (top) human donor pre-KD and post-KD fecal metagenomic and fecal metabolomic datasets and (bottom) mouse recipient pre-KD and post-KD fecal metagenomic, fecal metabolomic, serum metabolomic, hippocampal transcriptomic, and frontal cortical transcriptomic datasets. wKDA analyses was performed on the network. Red text denotes pathways, metabolites, or genes that were differentially regulated (p<0.05) between pre-KD and post-KD in prior individual dataset analyses (donor: n=10 patients per diet condition; recipient: n=10 per patient diet condition, where each sample is pooled from 5-6 recipient mice per donor patient sample). **(B)** Table of top associated genes from epilepsy GWAS mapping onto mouse recipient hippocampal and frontal cortical DEGs (n=10 per patient diet condition, where each sample is pooled from 6 recipient mice per donor patient sample). MMVEC, microbe-metabolite vectors; KD, ketogenic diet; wKDA, weighted key driver analysis; GWAS genome-wide association study; DEGs, differentially expressed genes

To gain insight into whether the hippocampal and frontal cortical transcripts that co-occur with microbial metagenomic and metabolomic features have been implicated in human epilepsy, single nucleotide polymorphisms (SNPs) identified from epilepsy genome-wide association studies (GWAS) were mapped to genes using hippocampus and frontal cortex splicing quantitative trait loci (sQTLs) and expression quantitative trait loci (eQTLs) to represent epilepsy-associated genes informed by GWAS. The mouse orthologs of these human GWAS disease genes were then compared with hippocampal and frontal cortical transcriptomic results to identify DEGs in post-KD vs pre-KD recipients that have been implicated in genetic risk for human epilepsy. There was a statistically significant enrichment of the hippocampal DEGs in the epilepsy GWAS (p=0.003), but no significant enrichment of the frontal cortical DEGs in the epilepsy GWAS (p=0.26) (**Figure 4B**). These results suggest that microbial alterations in hippocampal gene expression may contribute to the microbiome-dependent increases in seizure resistance seen in post-KD recipient mice compared to pre-KD controls. From the co-occurrence network, 5 hippocampal DEGs were linked to epilepsy GWAS results: *Atp5c*, which encodes mitochondrial ATP synthase; *Rprd2,* which encodes a transcriptional repressor that modulates RNA polymerase II activity; *Gnaz*, which encodes G protein alpha subunit that regulates ion equilibrium and chaperone-mediated folding; *Cfdp1*, which encodes a subunit of the chromatin remodeling complex and is important for cell division, and *Mro*, which encodes a nucleolus protein proposed to be testis-determining in the reproductive tract, but expressed in the nervous system with as yet unknown function. Overall, the multi’omics analysis of human donor and mouse recipient datasets together with epilepsy GWAS mapping to hippocampal and frontal cortical DEGs identified key microbial genomic pathways and microbially modulated metabolites that may contribute to alterations in the expression of particular hippocampal genes in mice that exhibit microbiome-induced protection against 6-Hz seizures.

## DISCUSSION

Results from this research provide evidence from a treatment study of children with refractory epilepsy, coupled with functional testing in gnotobiotic mice, that clinical KD regimens alter the function of the gut microbiome in ways that could contribute to seizure protection. We assessed microbiome composition and function in 10 children with refractory epilepsy shortly before initiating and approximately one month after adherence to classical KD regimens. Following clinical practice, the patient cohort was heterogeneous in type and underlying cause of refractory epilepsy, as well as the ratio of fat to carbohydrate and protein and specific nutritional composition of the KD they consumed (**Table S1**). This highlights the diversity of epilepsies that resist current antiepileptic drugs and the broad range of KD interventions that are administered to treat pediatric refractory epilepsy. Consistent with this heterogeneity, we observed that participants varied substantially in the composition of the fecal microbiota at baseline and in response to KD treatment. There was no clear KD-induced taxonomic signature of the gut microbiota that was shared across the study population, which contrasts prior studies of KD treatments for epilepsy that each reported alterations in the gut microbiota in response to a KD. Our results, however, support the finding that little to no consistency in specific microbial taxa affected exists across studies (Özcan et al., 2022).

Despite variation in microbiota composition, we observed evidence of shared functional features of the gut microbiome that were seen with KD treatment across participants in the study. This aligns with the notion of functional redundancy of gut microbes, wherein phylogenetically unrelated species can exhibit the same genetically-encoded biological activities (Tian et al., 2020). Results from metagenomic sequencing indicated that microbial genes related to fatty acid β-oxidation, 2-methylcitrate cycle, glycol metabolism, and proline biosynthesis were more highly represented in the gut microbiota of epileptic children after treatment with the KD compared to their internal pre-treatment controls. β-oxidation by select microbes in anaerobic environments enables them to utilize fatty acids from the diet as energy sources, wherein saturated and unsaturated fatty acids are oxidized into acetyl-CoA (Yao & Rock, 2017). β-oxidation of dietary odd-chain fatty acids additionally produces propionyl-CoA, which can be toxic to cells, so the methylcitrate cycle enables microbes to further catabolize propionyl-CoA into pyruvate and succinate (Dolan et al., 2018). Glycol, including glycolate and glyoxylate, metabolism allows microbes to use products from fatty acid oxidation to fuel gluconeogenesis (Ahn et al., 2016).

Proline synthesis from the central metabolite glutamate, via intermediates amino acids arginine and ornithine, is widely upregulated in bacteria to counteract growth in osmotically unfavorable conditions (Stecker et al., 2022). The elevated representation of genes related to these pathways in the post-KD samples suggests that the clinical KD shapes the gut microbiome to enrich microbial taxa that digest fat and synthesize carbohydrates under fat-rich, carbohydrate-limited conditions. These metagenomic features of the human microbiome from pediatric epilepsy patients consuming a clinical KD were preserved upon transfer to GF mice that were fed a standard diet, suggesting that the source microbes are maintained under non-ketogenic dietary conditions.

Aligning with results from metagenomic sequencing, metabolomic profiling of fecal samples from KD-treated epileptic children revealed statistically significant alterations in several metabolites, including subsets of amino acids, sugar acids, hydroxy fatty acids, bile acids, and other fatty acid derivatives, which reflect KD-, and potentially microbiome-, induced alterations in lipid and amino acid metabolism. In particular, glutamate and ornithine, both precursors of proline, were significantly decreased in post-KD human samples, relative to pre-KD controls, which may align with the observed metagenomic alterations in proline biosynthesis pathways. These metabolite alterations were induced by KD consumption in mice, and modified by microbiota depletion in mice, suggesting a causal response to the clinical KD in the human cohort that is dependent on the gut microbiome. Microbially modulated increases in palmitoleoylcarnitine (C16:1) were also seen in KD-fed mice and in post-KD human samples, alongside several other lipid species, aligning with the high fat content of the KD and roles for the gut microbiome in lipid metabolism (Joyce et al., 2014).

The individual metabolite changes seen in human donors, including those induced by KD in a microbiome-dependent manner, were not specifically recapitulated by microbiome transfer to GF mice that were fed standard chow. This is perhaps not surprising given the important role of dietary composition in driving microbial activity (David et al., 2014). While no specific metabolite shifts were shared, a few pathway-level metabolomic changes were consistent between post-KD fecal samples from human donor (consuming the clinical KD) and mouse recipients (fed standard chow), relative to their respective pre-KD controls. Namely, differentially abundant metabolites related to metabolism of methionine, glycine, serine, and betaine were shared across post-KD conditions for human donor and microbiota-recipient mice. Methionine metabolism involves the production of homocysteine, adenosine, cysteine, and alpha-ketobutyrate, which can then be routed to glucogenic pathways by conversion to propionyl- and succinyl-CoA. Serine, synthesized via glycerate, is used to create glycine (and cysteine) via the homocysteine cycle, which can undergo microbial conversion into pyruvate or glyoxylate. Betaine (trimethylglycine), derived from diet or synthesized from choline, is metabolized by the gut microbiome (Koistinen et al., 2019) and functions as a methyl donor in transmethylation reactions, including those involved in methionine metabolism. While the relevance to KD and seizure protection is unclear, alterations in peripheral and central amino acid metabolism have been widely implicated in mediating the anti-seizure effects of the KD (Yudkoff et al., 2001). In addition, post-KD samples from both human donors and mouse recipients exhibited alterations in chemicals related to lipid metabolism, such as hydroxy fatty acids. The shared metabolite pathway- and chemical subclass-level features may reflect changes that are induced by KD in humans and generally phenocopied by gut microbes upon transfer to mice reared under non-ketogenic conditions. This suggests that transfer of clinical KD-induced gut microbes to mice maintained under non-ketogenic conditions could result in molecular outputs that are distinct, but functionally similar, to those seen in the donor human sample.

We observed that inoculating mice with human fecal samples collected after clinical KD treatment conferred resistance to 6-Hz seizures compared to controls that received the baseline pre-treatment (pre-KD) microbiota. There was no correlation with patient responsiveness to diet, as indicated in clinician notes taken at 1 month after adherence to the clinical KD. This may be due to the unreliability of the metric, which was based on parental reporting, as well as the cross-sectional nature of the assessment, given inter-individual variation in latency to respond to KD treatments and the patient’s peak KD ratio. These concerns aside, the results highlight the importance of host determinants of KD responsiveness, some of which may mask or block any beneficial influences of the KD-associated microbiota. Many patients included in this study exhibited genetic bases for refractory epilepsy, some of which could be epistatic to functional genomic changes in the KD-associated gut microbiome. Large human studies that subclassify different types of epilepsies and seizure semiologies are warranted to study potential roles for the gut microbiome in modifying or predicting responsiveness to the KD.

The microbiota-dependent increases in seizure protection were associated with brain transcriptomic alterations. In particular, both hippocampus and frontal cortex from post-KD recipient mice exhibited enrichment of differentially expressed genes related to i) RNA processing, transcriptional regulation, and translation ii) TORC1 signaling and cell cycle, and iii) oxidative phosphorylation and cellular stress response (i.e., nitric oxide, reactive oxygen species), when compared to controls colonized with pre-KD microbiota from both GO enrichment and STRING network analyses. Neuronal excitability requires protein synthesis in response to altered neuronal stimulation, and risk factors for various epilepsies include dysregulation of RNA processing, RNA stability, transcription, and translation (Malone & Kaczmarek, 2022). TORC1 is a major nutrient-and energy-sensing serine/threonine kinase complex that controls cell growth and differentiation by coordinating core processes of transcription, translation, and autophagy. Abnormal regulation of TORC1 signaling has been implicated in a wide variety of epilepsies, and as such, is a therapeutic target of interest for treating refractory epilepsies (Nguyen & Bordey, 2021). Previous studies have reported inhibitory effects of the KD and select fatty acids on TORC1 activity (McDaniel et al., 2011; Warren et al., 2020), suggesting that it may contribute to the anti-seizure effects of the KD. Oxidative phosphorylation is a central process for cellular energy metabolism from nutrients, that generates as a byproduct reactive oxygen species (ROS) (Rowley & Patel, 2013) and is regulated by the retrograde glutamatergic neurotransmitter nitric oxide (NO). In animal epilepsy models, both ROS and NO are elevated during seizure activity due to oxidative stress-associated neuronal death (Zhu et al., 2017), which can further contribute to epileptogenesis (Chan, 2001). The KD has been previously reported to reduce oxidative stress by promoting antioxidant enzymatic activity and scavenging ROS (Greco et al., 2016). Overall, these results suggest that the KD-associated human gut microbiota alters brain transcriptional pathways that may contribute to protection against 6-Hz seizures in mice.

Integration of multi’omics datasets across human donor and mouse recipients revealed network associations between select gut microbial metagenomic pathways, fecal metabolites, serum metabolites, and hippocampal transcripts, suggesting that they may contribute to the microbiome-dependent increases in seizure protection seen in mice inoculated with human post-KD microbiota, compared to pre-KD controls. The human donor and mouse recipient co-occurrence networks were linked by shared metagenomic pathway nodes related to BCAA biosynthesis, CoA biosynthesis, L-alanine fermentation, and L-arginine biosynthesis. Key drivers for BCAA, CoA, and L-arginine biosynthesis were linked to hippocampal transcript modules enriched for genes related to neurogenesis and Wnt signaling. BCAAs modulate brain import of precursors required for synthesis of monoamine transmitters (Larsson & Markus, 2017; Salcedo et al., 2021; Song et al., 2017). BCAAs also serve as nitrogen donors for synthesis of glutamate vs. GABA, and as such regulates synaptic balance between excitation and inhibition, a key determinant of seizure susceptibility (McKenna et al., 2019). Wnt signaling regulates calcium pathways that are important for hippocampal neurogenesis and dendrite formation, and is increasingly linked to early epileptogenesis (Hodges and Lugo, 2018). Additionally, mapping GWAS-based risk genes to the co-occurrence network identified five hippocampal nodes as linked to epilepsy. Of particular interest was *Gnaz*, which encodes G protein alpha-Z, a protein that mediates neuronal signal transduction within the hippocampus (Jang et al., 2018) and is proposed to modulate seizure susceptibility (Hultman et al., 2019). BCAA derivatives have been reported to promote phosphorylation of G-proteins, and abnormalities in GPCR mediated neuronal signaling can contribute to increased susceptibility to seizure (Shellhammer et al., 2017; Yu et al., 2019). Altogether, results from this study reveal that the clinical KD regimens used to treat pediatric refractory epilepsy are associated with alterations in the function of the child microbiome, which causally modify brain function and seizure susceptibility upon transfer to mice. Further research is warranted to define the mechanisms by which the human KD-associated microbiome signals across the gut-brain axis to modify seizure risk, and to further assess the potential for identifying microbiome-based interventions that could increase the efficacy of KD treatment, alleviate dietary side effects, and/or ease clinical implementation.

## LIMITATIONS OF STUDY

A key limitation of this study design is the prioritization of experimental reproducibility, which included cohorts of 14-16 mice per patient sample, over patient sample size, which included 10 children with refractory epilepsy, each sampled before and at approximately 1 month after adherence to a clinical KD. We reasoned that by internally controlling for baseline microbiota for each patient, we could effectively evaluate microbial alterations in response to the clinical KD within a relatively small patient group. We further posited that this study design would enable us to sample from a heterogenous patient population reflective of the etiopathological variation typically seen in refractory epilepsy. It would also us to be inclusive of the wide range of individuals with pediatric epilepsy who typically seek clinical KD treatment. This level of diversity in a small patient population may have contributed to our finding that there was no shared taxonomic response of the gut microbiome to the clinical KD, despite some shared functional genomic features when considering all post-KD microbiota relative to all pre-KD controls.

Additional constraints of the study, as discussed in the main text, are the inherent technical and biological shortcomings in “transplanting” microbiota across different mammalian hosts. In this study, we achieved levels of human-to-mouse microbiota “transplant” fidelity analogous to those reported in existing literature even when we maintained mice on a conventional rather than ketogenic diet (Bokoliya et al., 2021; Kennedy et al., 2018; Walter et al., 2020). However, the discrepancies between recipient and donor microbiota draw into question the relevance of findings in gnotobiotic mice to the human condition. To help mitigate this, we focused entirely on features of the gut microbiome that were differential between post-KD and pre-KD conditions and shared between human donors and mouse recipients. However, we acknowledge that artifacts of the microbiota transfer approach, which are not relevant to the clinical condition, may contribute to the microbiome-dependent functional differences observed in the gnotobiotic mouse experiments in this study. Nevertheless, the observed results provide important proof-of-principle that differences in the function of the gut microbiota regulate seizure susceptibility.

In assessing causal relationships between the KD-associated microbiome and host physiologies linked to seizure susceptibility, we made the major assumption that there exists a singular microbiome-dependent mechanism to increase seizure threshold that was common across all post-KD mouse recipient cohorts relative to all pre-KD cohort controls. Our analysis does not take into account the possibility that there are multiple microbiome-dependent mechanisms that are distinct and that each result in resistance to 6-Hz seizures. Expanded studies that involve subclassification of the human participants and/or mouse recipients would aid in addressing this prospect.

Moreover, we chose to study the 6-Hz psychomotor seizure model based on its widespread use as a model of refractory epilepsy (Kehne et al., 2017), its utility for screening novel antiepileptic drugs (Barton et al., 2001), and its responsiveness to the KD (Hartman et al., 2008). We also reasoned that its measure of acutely induced seizures would preclude confounding effects of chronic genetic mutation or kindling-based models on modifying the gut microbiome (Löscher, 2017). Further research is needed to assess roles for the KD-induced gut microbiome in modifying seizures across additional epilepsy models to determine whether particular seizure semiologies or types of epilepsy are more amenable to modulation by the gut microbiome.

In light of the aforementioned heterogeneity in patient population, small sample size, variation in clinically-guided KD regimens, discrepancies introduced by the microbiota “transplantation” approach, cross-species and diet comparisons (i.e., human on KD, mouse on standard diet), and assumptions adopted for data analysis, our statistical analyses for shotgun metagenomic, untargeted metabolomic, and bulk transcriptomic data were performed with lenient thresholds for differential abundance (p<0.05), with a focus on pathway-level signatures that were dependent upon the clinical KD and conferred by the KD-associated microbiota. Notably, for all animal experiments, technical replicates per donor sample were averaged, and only the biological (i.e., donor) N was used for statistical analysis. Despite the expected variability, we detected consistent KD-dependent alterations in microbial genes and metabolites in epileptic children undergoing dietary treatment, and further observed KD- and microbiome-dependent alterations in metagenomic pathways and metabolomic pathways that were shared across human donor and microbiome-recipient mice when using these parameters. Brain transcriptomic signatures were seen when comparing all mouse recipient cohorts receiving the post-KD microbiome (all of which exhibited resistance to 6-Hz seizures) relative to those receiving the pre-KD controls. Finally, results for all seizure testing experiments, which revealed shared phenotypic outcomes for post-KD groups compared to pre-KD groups, were well-powered and analyzed according to conventional statistical methods (Festing & Altman, 2002). All caveats considered, the results from this study extend existing pre-clinical research to provide initial evidence that clinical KD treatments shape the function of the gut microbiome of children with refractory epilepsy in ways that have the potential to causally modify seizure susceptibility. Continued research is warranted to elucidate the particular microbial functional activities that act together to modify signaling across the gut-brain axis to promote seizure protection and to further assess the potential to apply microbiome-based interventions to treat refractory epilepsy.

## ACKNOWLEDGEMENTS

We thank members of the Hsiao laboratory for their critical review of the manuscript, and members of the UCLA Goodman Luskin Microbiome Center Gnotobiotics Core Facility for technical support. This work was supported by funds from a UCLA Whitcome Fellowship to G.R.L. and National Institute of Neurological Disorders and Stroke (NINDS) grant (#R01 NS115537) to E.Y.H. E.Y.H. is a New York Stem Cell Foundation – Robertson Investigator. Individuals involved in this research were supported in part by the New York Stem Cell Foundation and grant number 2018- 191860 from the Chan Zuckerberg Initiative DAF, an advised fund of Silicon Valley Community Foundation. Brain RNA sequencing and a subset of metagenomic sequencing was supported by funds from Bloom Science, Inc. (see Declaration of Interests). None of the funding sources influenced or provided input on data analysis or interpretation.

## AUTHOR CONTRIBUTIONS

G.R.L., S.M.H, C.A.O., M.B., and J.P. performed the experiments and analyzed the data, B.R., and J.H.M. led the clinical study. G.R.L., C.A.O., J.H.M., X.Y., and E.Y.H. designed the study,

G.R.L. and E.Y.H. wrote the manuscript. All authors discussed the results and commented on the manuscript.

## DIVERSITY AND INCLUSION

We worked to ensure sex balance in the selection of human subjects. One or more of the authors of this paper self-identifies as an underrepresented ethnic minority in science. While citing references scientifically relevant for this work, we also actively worked to promote gender balance in our reference list.

## STAR★METHODS

Detailed methods are provided in the online version of the paper and include the following:

- KEY RESOURCES TABLE
- RESOURCE AVAILABILITY
  - Lead Contact
  - Materials availability
  - Data and code availability
- EXPERIMENTAL MODELS AND SUBJECT DETAILS - Human Fecal Samples
- Mice
- METHOD DETAILS - 16S rRNA Gene Sequencing
- Fecal Shotgun Metagenomics
- Human Donor Fecal Microbiota Transfer
- 6-Hz Psychomotor Seizure Assay
- Antibiotic Treatment
- Fecal and Serum Metabolomics
- Transcriptomics
- Multi’omics Integration
- Marker set enrichment analysis (MSEA) to connect hippocampus and frontal cortex DEGs with epilepsy GWAS
- QUANTIFICATION AND STATISTICAL ANALYSIS

## KEY RESOURCES TABLE

| REAGENT or RESOURCE | SOURCE | IDENTIFIER |
| --- | --- | --- |
| **Biological samples** |  |  |
| Stool samples from human subjects | This study | N/A |
| **Chemicals, Peptides, and Recombinant Proteins** |  |  |
| Vancomycin hydrochloride | Chem-Impex International | 00315 |
| Neomycin trisulfate salt hydrate | Sigma-Aldrich | N1876 |
| Metronidazole | Sigma-Aldrich | M1547 |
| Ampicillin sodium salt | Sigma-Aldrich | A9518 |
| TURBO DNase | Invitrogen | AM2238 |
| Ultrapure water | ThermoFisher | 10977015 |
| 1x PBS | ThermoFisher | 10010023 |
| Tetracaine Hydrochloride Ophthalmic Solution, USP 0.5% | Oceanside Pharmaceuticals | 68682-920-64 |

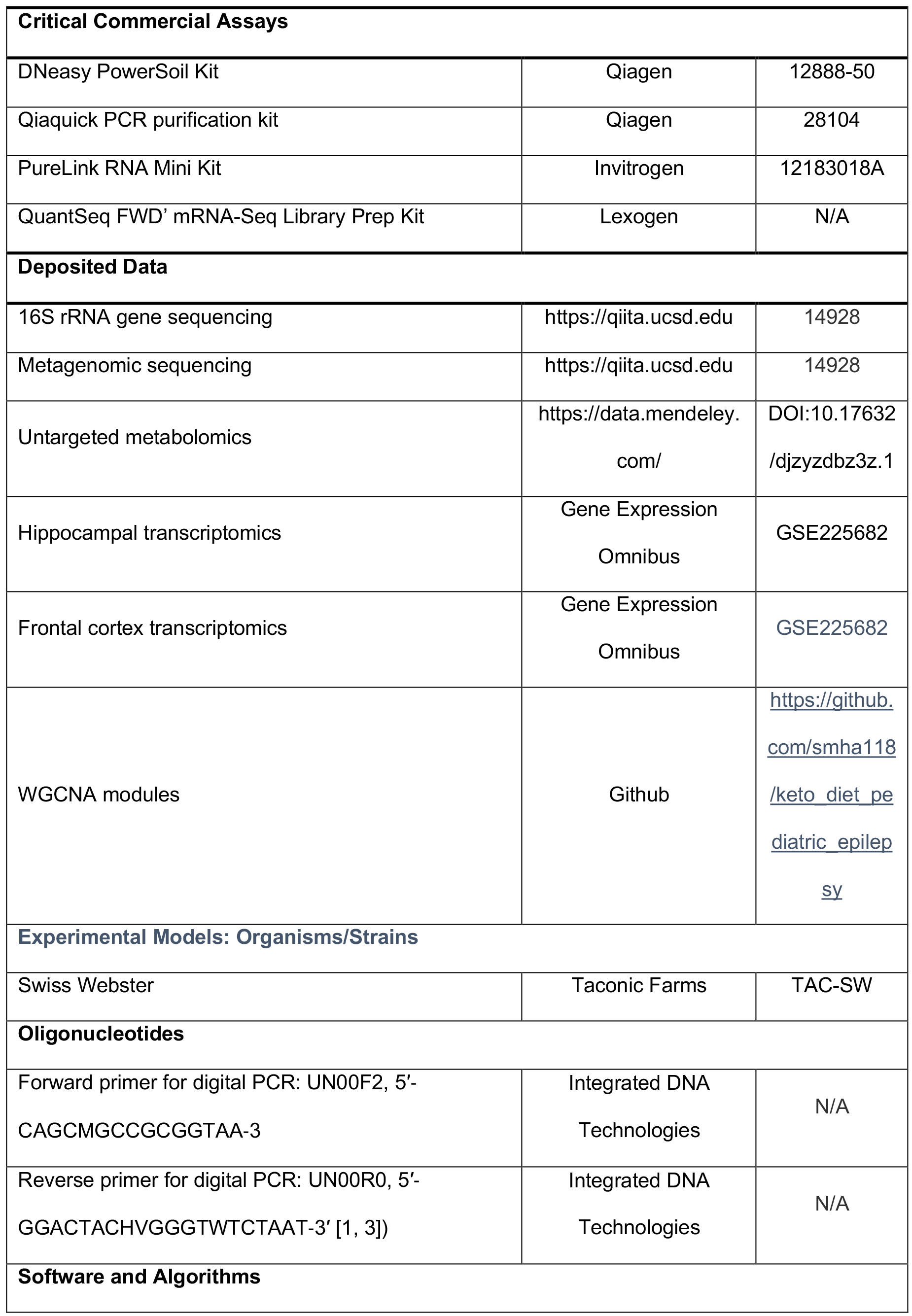

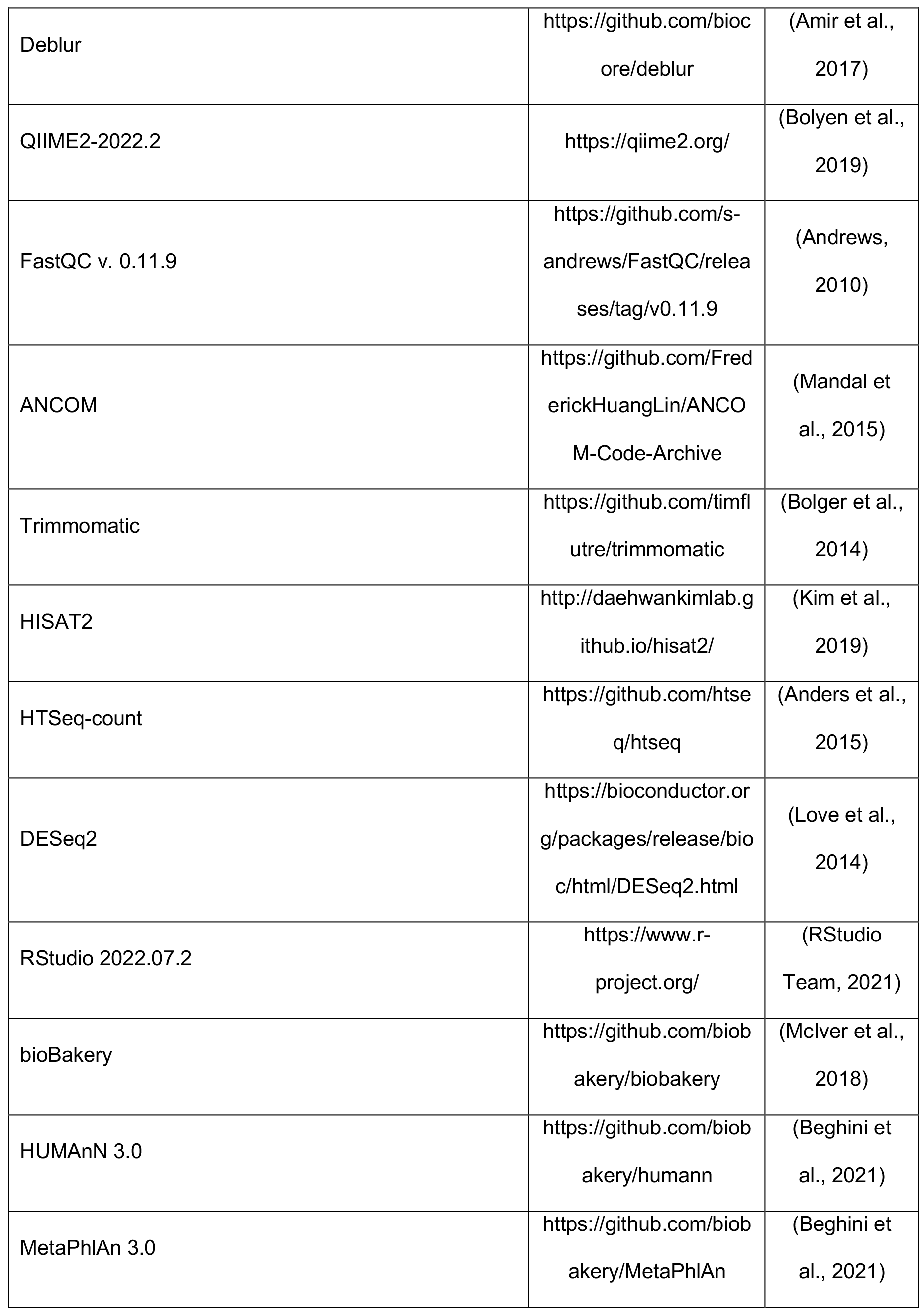

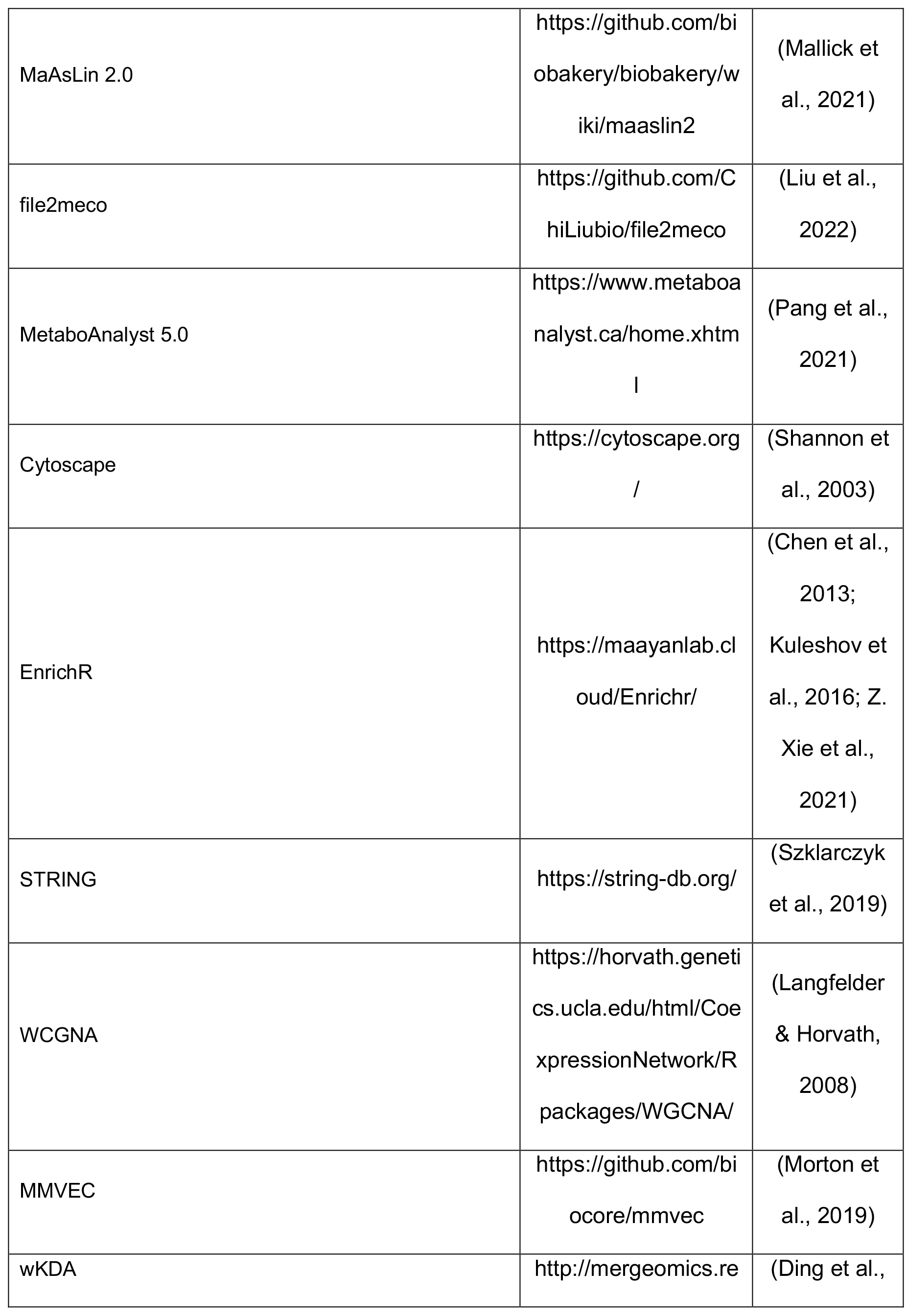

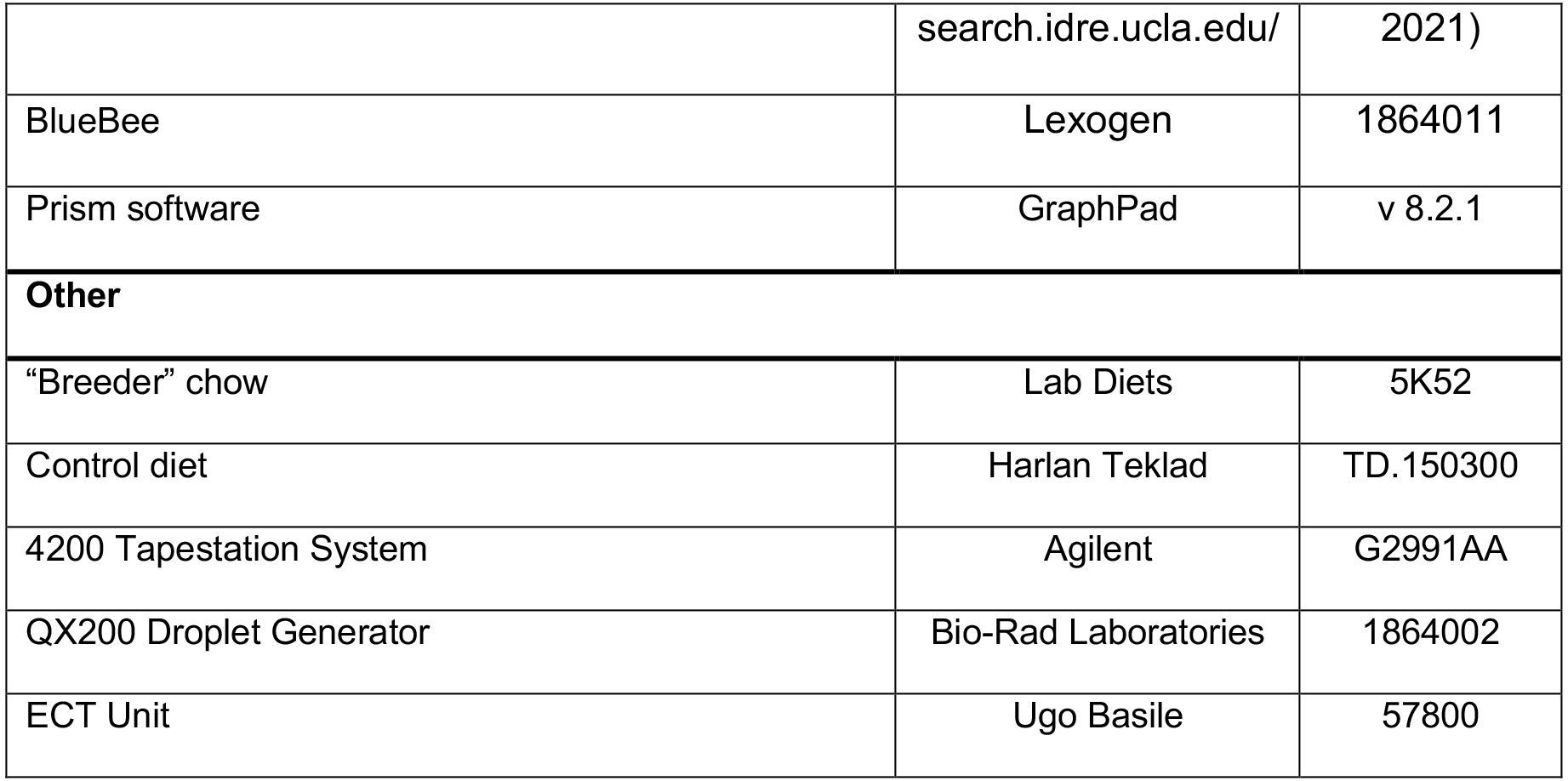

## RESOURCE AVAILABILITY

### Lead Contact

Further information and requests for resources and reagents should be directed to and will be fulfilled by the Lead Contact, Elaine Hsiao

### Materials Availability

This study did not generate new unique reagents.

### Data and Code Availability

Data from 16S rRNA gene sequencing, metagenomic profiling, and associated metadata are presented in Tables S2, S3, S5, and S6 are available online through the QIITA repository (https://qiita.ucsd.edu/) with the study accession #14928. Metabolomic data are presented in Tables S7, S8, S9, and S10 and are available online through Mendeley data with DOI:10.17632/djzyzdbz3z.1. Transcriptomic data are presented in Tables S11 and S12 and available online through Gene Expression Omnibus repository with the identification number #GSE225682.

## EXPERIMENTAL MODELS AND SUBJECT DETAILS

### Human Subjects

This study was approved by UCLA’s Institutional Review Board (IRB protocol #15-000453). Pediatric refractory epilepsy patients were screened and enrolled in collaboration with the Ketogenic Diet Program at UCLA Mattel Children’s Hospital. Prospective participants who met study criteria were provided information detailing this study by phone and email 1-2 weeks before their pre-diet initiation visit. Prior to enrollment, informed signed consent was provided by all participants and their guardians to the program clinical coordinator during the pre-diet initiation appointment. Subjects were enrolled across diverse seizure semiology and prior medical histories. Inclusion criteria: enrolled in UCLA’s program for classical 4:1 KD, children aged 1-10 with refractory epilepsy, any gender, any ethnicity, any previous exposure to AEDs, any seizure semiology. Exclusion criteria: use of antibiotics or probiotics within 7 days prior to enrollment, existing diagnosis of gastrointestinal, immunological, or metabolic disorder. Human donor stool samples were collected from 10 participants, each providing 2 stool samples. The first sample was collected within 1 day before starting KD treatment (pre-KD) and the second sample was collected after maintaining on the clinical KD for 1 month (post-KD). Clinical metadata from the medical record were coded and stripped of identifiers before being shared, and included participant demographic data, medical history, AED exposure history, additional medications take during this study, laboratory blood glucose and bloody ketone body levels, seizure severity, seizure frequency, seizure semiology, and dietary regimen (Table S1).

### Human stool sample collection

For in-patient fecal sample collection, once a study participant was admitted to the hospital during the pre-diet initiation visit, they were given a coded stool collection kit and sterile specimen container. Stool samples were freshly collected within 1 day prior to starting the clinical KD treatment (pre-KD). Fresh stool samples were immediately placed on dry ice for short term storage and transportation and were freshly frozen at −80°C for long-term storage. Post-KD stool samples were collected in the same manner as stated above when the study participant returned for the 1-month follow-up visit. For out-patient collection of the post-KD stool sample, which was necessitated because of hospital pandemic policies, a deidentified stool sample collection kit and sterile specimen cup was provided to the patient and guardian along with a pre-labeled return shipping box. After 1 month of the clinical KD treatment, stool samples were collected in a sterile specimen cup, immediately placed in an at home freezer, and the next day either (1) shipped back overnight to UCLA on dry ice or (2) brought with the patient to their 1-month follow-up appointment. Fresh frozen fecal samples were homogenized under liquid nitrogen and 3 ∼500 mg aliquots were made per sample by sterile storage in anaerobic Balch tubes to be used for transplantation, metagenomic, and metabolomic studies.

### Mice

6-8 week old wild-type germ-free Swiss Webster mice (Taconic Farms), were bred in UCLA’s Center for Health Sciences Barrier Facility. Breeding animals were fed ‘‘breeder’’ chow (Lab Diets 5K52). Experimental animals were fed vitamin- and mineral-matched control diet (Harlan Teklad TD.150300). Juvenile mice were used to mimic the age range of the human donor population (<10 years old). All animal experiments were approved by the UCLA Animal Care and Use Committee.

## METHOD DETAILS

### 16S rRNA Gene Sequencing and Analysis

Bacterial genomic DNA was extracted from human or mouse fecal samples using the Qiagen PowerSoil Kit. For human samples, the n reflects one donor sample. For mouse samples, the n reflects independent cages containing 3 mice per cage to preclude effects of co-housing on microbiota composition. The sequencing library was generated in line with (Caporaso et al., 2011). PCR amplification, run in triplicate, of the V4 region of the 16S rRNA gene was completed using individually barcoded universal primers and 30 ng of the extracted genomic DNA. The PCR product triplicates were pooled and purified using the Qiaquick PCR purification kit (Qiagen). Samples were sequenced using the Illumina MiSeq platform and 2 x 250bp reagent kit for paired-end sequencing at Laragen, Inc. Amplicon sequence variants (ASVs) were chosen by closed reference clustering based on 99% sequence similarity to the SILVA138 database. Taxonomy assignment, rarefaction, and differential abundance testing were performed using QIIME2 2022.2 (Bolyen et al., 2019; Mandal et al., 2015).

### Fecal Shotgun Metagenomics

Bacterial genomic DNA was extracted from human or mouse fecal samples using the Qiagen PowerSoil Kit. 1 ng of DNA was used to prepare DNA libraries using the Nextera XT DNA Library Preparation Kit (Illumina) and genomic DNA was fragmented with Illumina Nextera XT fragmentation enzyme. IDT Unique Dual Indexes were added to each sample before 12 cycles of PCR amplification. AMpure magnetic Beads (Beckman Coulter) were used to purify DNA libraries which were eluted in QIAGEN EB buffer. Qubit 4 fluorometer and Qubit dsDNA HS Assay Kit were used for DNA library quantification. Libraries were then sequenced on Illumina HiSeq 4000 platform 2×150bp at a 6M read depth using by CosmosID. Metagenomic data was analyzed using HUMAnN 3.0 (Beghini et al., 2021) and MetaCyc database to profile gene families and pathway abundance. File2meco R package was used for MetaCyc pathway hierarchical classification (Liu et al., 2022). MaAsLin 2.0 (Mallick et al., 2021) was used to assess significant pathway associations between pre-KD and post-KD with a p-value cutoff of 0.1, where p < 0.05 pathways are indicated in the figure by asterisk. Heatmaps were generated using the pheatmap v1.0.12 package for R.

### Human Donor Fecal Microbiota Transfer

To prepare collected human stool samples for transplantation studies, the frozen stool sample was pulverized into a powder under liquid nitrogen stream in a sterile heavy-duty foil covered mortar and pestle, aliquoted at 500 mg per tube into 2mL screw cap tubes, and frozen at −80C. A single 500 mg aliquot of human stool sample was entered into a Coy anaerobic chamber and resuspended in pre-reduced 1x PBS + 0.05% L-cysteine. The sample was homogenized using sterile borosilicate glass beads and passed through a 100um filter. GF Swiss Webster mice were colonized by oral gavage of 200 ul fecal suspension. Excess fecal suspension was resuspended and stored at −80C in pre-reduced 1x PBS + 0.05% L-cysteine + 15% glycerol. For administration of fecal filtrates, the fecal suspension was passed through a sterile 0.2 um filter before colonization via oral gavage using 200 ul fecal filtrate.

### 6-Hz Psychomotor Seizure Assay

6-Hz psychomotor seizure assay testing was conducted following Samala et al., 2008. One drop (∼50 ul) of 0.5% tetracaine hydrochloride ophthalmic solution was applied to the corneas of each mouse 15 min before stimulation. A thin layer of electrode gel (Parker Signagel) was applied directly to the corneal electrodes and was reapplied before each trial. A constant-current current device (ECT Unit 57800, Ugo Basile) was used to deliver current through the corneal electrodes at 3s duration, 0.2 ms pulse-width and 6 pulses/s frequency. CC50 (the milliamp intensity of current required to elicit seizures in 50% of the mouse cohort) was measured as a metric for seizure susceptibility. Pilot experiments were conducted to identify 28 mA as the CC50 for SPF wild-type Swiss Webster mice, aged 6-8 weeks. Each mouse was seizure-tested only once, and thus at least n > 14 mice were used to adequately power each cohort. To determine CC50s for each tested cohort, 28 mA of current was administered to the first mouse per cohort, followed by stepwise fixed increases or decreases by 2 mA intervals. Mice were restrained manually during stimulation and then released into a new cage for behavioral observation. Quantitative measures for falling, tail dorsiflexion (Straub tail), forelimb clonus, eye/vibrissae twitching and behavioral remission were scored manually. For each behavioral parameter, we observed no correlation between percentage incidence during 28+ mA seizures between pre-KD or post-KD microbiota status, suggesting a primary effect of the microbiota on seizure incidence rather than presentation or form. Latency to exploration (time elapsed from when an experimental mouse is released into the observation cage (after corneal stimulation) to its first lateral movement) was scored manually with an electronic timer. Mice were blindly scored as protected from seizures if they did not show seizure behavior and resumed normal exploratory behavior within 10 s. Seizure threshold (CC50) was determined as previously described (Kimball et al., 1957), using the average log interval of current steps per experimental group, where sample n is defined as the subset of animals displaying the less frequent seizure behavior. Data used to calculate CC50 are also displayed as latency to explore for each current intensity, where n represents the total number of biological replicates per group regardless of seizure outcome.

### Antibiotic Treatment

Transplanted mice were gavaged with a solution of vancomycin (50 mg/kg), neomycin (100 mg/kg) and metronidazole (100 mg/kg) every 12 hours daily for 5 days, as adapted from (Reikvam et al., 2011). Ampicillin (1 mg/ml) was provided *ad libitum* in drinking water. For mock treatment, mice were gavaged with a similar volume of 1x PBS (vehicle) water every 12 hours daily for 7 days. Antibiotic-treated mice were maintained in sterile caging with sterile food and water and handled aseptically for the remainder of the experiments.

### Fecal and Serum Metabolomics

Previously collected human donor fecal samples were aliquoted as described in section “Human Donor Fecal Microbiota Transfer”. Mouse fecal samples were collected from mice housed across independent cages, with four cages housing 3 mice and one cage housing 2 mice. Mouse serum samples were collected by cardiac puncture and separated using SST vacutainer tubes, then frozen at −80C. Samples were prepared using the automated MicroLab STAR system (Hamilton Company) and analyzed on GC/MS, LC/MS and LC/MS/MS platforms by Metabolon, Inc. Protein fractions were removed by serial extractions with organic aqueous solvents, concentrated using a TurboVap system (Zymark) and vacuum dried. For LC/MS and LC-MS/MS, samples were reconstituted in acidic or basic LC-compatible solvents containing > 11 injection standards and run on a Waters ACQUITY UPLC and Thermo-Finnigan LTQ mass spectrometer, with a linear ion-trap frontend and a Fourier transform ion cyclotron resonance mass spectrometer back-end. For GC/MS, samples were derivatized under dried nitrogen using bistrimethyl-silyl-trifluoroacetamide and analyzed on a Thermo-Finnigan Trace DSQ fast-scanning single-quadrupole mass spectrometer using electron impact ionization. Chemical entities were identified by comparison to metabolomic library entries of purified standards. Following log transformation and imputation with minimum observed values for each compound, post-KD vs. pre-KD comparisons for human fecal, and mouse serum and fecal data were analyzed by paired t-test. Metabolomic data from SPF or antibiotic-treated mice fed KD vs. CD chow were acquired from (Olson et al., 2018), as log transformed and imputed with minimum observed values for each compound. Data were analyzed using two-way ANOVA to test for group effects. P and q-values were calculated based on two-way ANOVA contrasts. Principal components analysis was used to visualize variance distributions. Supervised Random Forest analysis was conducted to identify metabolomics prediction accuracies. Metabolite set enrichment analysis (MSEA) using the Metaboanalyst 5.0 platform (Pang et al., 2021) was performed on human fecal, mouse fecal, and mouse serum metabolites statistically significantly altered in post-KD compared to pre-KD (p-val<0.05). Metabolite sets were analyzed for chemical sub-class enrichment and metabolite pathway enrichment, using The Small Molecule Pathway Database (SMPDB).

### Transcriptomics

Recipient mice were sacrificed on day 4 post-colonization. Hippocampal, frontal cortex, were dissected from pre-KD and post-KD recipient mice (n=6 per cohort) and immediately placed in Trizol. RNA was extracted using the PureLink RNA Mini kit with Turbo DNAse treatment. RNA was prepared using the TruSeq RNA Library Prep kit and 2 Å∼ 69-bp paired-end sequencing was performed using the Illumina HiSeq 4000 platform by the UCLA Neuroscience Genomics Core. FastQC v0.11.5, bbduk v35.92, and RSeQC v2.6.4 were used for quality filtering, trimming, and mapping. Reads were aligned to UCSC Genome Browser assembly ID: mm10 using STAR v2.5.2a, indexed using samtools v1.3, and aligned using HTSeq-count v0.6.0. Differential expression analysis was conducted using DESeq2 v1.24.041. Heatmaps were generated using the pheatmap v1.0.12 package for R. GO term enrichment analysis of differentially expressed genes with p < 0.05 was conducted using enrichR v3.1. Protein interaction networks were generated using STRING v10.5. Functional enrichments of network nodes were categorized by GO: biological process, molecular function, and cellular component.

### Multi-omics Integration

To assess the relationships across omics layers, we first carried out dimension reduction for each data set using weighted gene co-expression network analysis (WGCNA v1.72.1) (Langfelder & Horvath, 2008). Metabolomics for human donors and mouse recipients and RNA-seq for mouse recipients (hippocampus and frontal cortex) were used to build WGCNA modules within each dataset, where modules represent clusters of highly co-regulated/expressed molecules which are typically involved in similar biological functions. For metabolomics data, *goodSamplesGenes* function was first used with default parameters to filter out sparse metabolites across samples before constructing networks; this step was not used for RNAseq data. Standard WGCNA steps were then carried out for the filtered metabolomics and RNAseq data. Module eigengenes (MEs), or the first axis of principal component were calculated from each module. MEs were then targeted for correlation analysis with the metadata (pre-KD vs. post-KD and responder vs. non-responder). Modules that had significant correlation (p-val <0.05) with the metadata were chosen for subsequent integrative analysis.

A systematic network that combined all omics data was inferred based on the probability of co-occurrence (POC) between molecules from different omics data. To calculate POC, we leveraged a neural-net based tool called MMVEC v1.0.6 with default parameters (Morton et al., 2019). The subset of raw data that contains module components that were selected from WGCNA analysis were log normalized and combined based on sample ID. This combined data matrix was then used as input for MMVEC. For example, on donor side, modules from fecal metagenome and metabolomics were added together and, on the recipient side, the combined matrix contained the raw data from metagenome, metabolomics, and RNAseq. Due to high density of the overall network generated from MMVEC, the top 10% of POC connections were retrieved to minimize overall complexity of the network for both donors and recipients using in-house python script (https://github.com/smha118/keto_diet_pediatric_epilepsy).

The networks of modules from individual omics layers from donor (metagenome, metabolomics) and recipient (metagenome, metabolomics, RNAseq) as well as differentially expressed/abundance molecules were then seeded into Mergeomics v3.16 pipeline along with the integrated network generated with MMVEC for weighted key driver analysis (wKDA) to identify key drivers of the networks (Ding et al., 2021). wKDA uses a χ2 -like statistic to identify molecules that are connected to significant larger module components than what would be expected by random chance. The analysis was done on the human and mouse networks separately. To further look into the network that are relevant to ketogenic diet and epilepsy, we selected key drivers (KDs) based on i) the number of modules that a key driver was invoked related to, ii) their relation to the Ketogenic diet or epilepsy. A subset of nodes in each module that were connected to the KDs were collected. These nodes were retrieved with following priority i) they are part of differentially regulated molecules ii) POC value with KDs. Finally, the network was visualized using Cytoscape (Shannon et al., 2003). To minimize overall density of the network, we chose to show the key drivers Mergeomics with the highest occurrence in their respective MEs and with > 5 degrees of connectivity.

### Marker set enrichment analysis (MSEA) to connect hippocampus and frontal cortex DEGs with epilepsy GWAS

To assess the potential role of the DEGs from the hippocampus and frontal cortex in epilepsy, we collected the summary statistics of the latest epilepsy GWAS (Abou-Khalil et al., 2018). Single nucleotide polymorphisms (SNPs) that had a linkage disequilibrium of r^2^> 0.5 were filtered to remove redundancies. To map the epilepsy GWAS SNPs to genes, we used GTEx version 8 eQTL and sQTL data for brain hippocampus and brain frontal cortex (Aguet et al., 2020), which help us derive genes likely to be regulated by the SNPs. We next used the MSEA function of the Mergeomics package (Ding et al., 2021) to compare epilepsy disease association p-values of the SNPs representing the DEGs (hippocampus or frontal cortex) with those of the SNPs mapped to random genes to assess whether the DEGs contain SNPs that show stronger epilepsy association than random genes using a χ^2^-like statistic.

## QUANTIFICATION AND STATISTICAL ANALYSIS

Statistical analyses were conducted using Prism8 software v8.2.1 (GraphPad). Before statistical analysis, data was assessed for distribution to determine appropriate statistical tests to use. Data were plotted in figures as mean ± SEM. For figures: 1B, S2C, S3B, S3B, S4C, *n =* the number of technical replicates. For all other figures, *n =* the number of biological replicates. No samples or animals were excluded from data analysis. Differences between two sample conditions from parametric data sets were analyzed using two-tailed, paired Student’s t-test. Differences between two sample conditions from nonparametric data sets were analyzed using two-tailed, Wilcoxon matched-pairs signed rank test. For differences among >2 groups when analyzing one variable, a one-way ANOVA with Tukey’s post hoc test was used. For differences among ≥2 groups with two variables, a two-way ANOVA with Sidak’s post hoc test was used. For technical replicates from within-patient analysis (Figures: 1B, S2C, S3B, S3B, S4C), differences from the above tests are denoted by: ^#^p<0.05; ^##^p<0.01; ^###^p<0.001; ^####^p<0.0001. For biological replicates (all other figures), differences from the above tests are denoted by: *p<0.05; **p<0.01; ***p<0.001; ****p<0.0001. Non-significant differences are denoted in the figures using “n.s”.

## SUPPLEMENTAL INFORMATION

Supplemental Information includes seven figures and fourteen tables that contain source data and can be found with this article.

---

**Figure S1:**
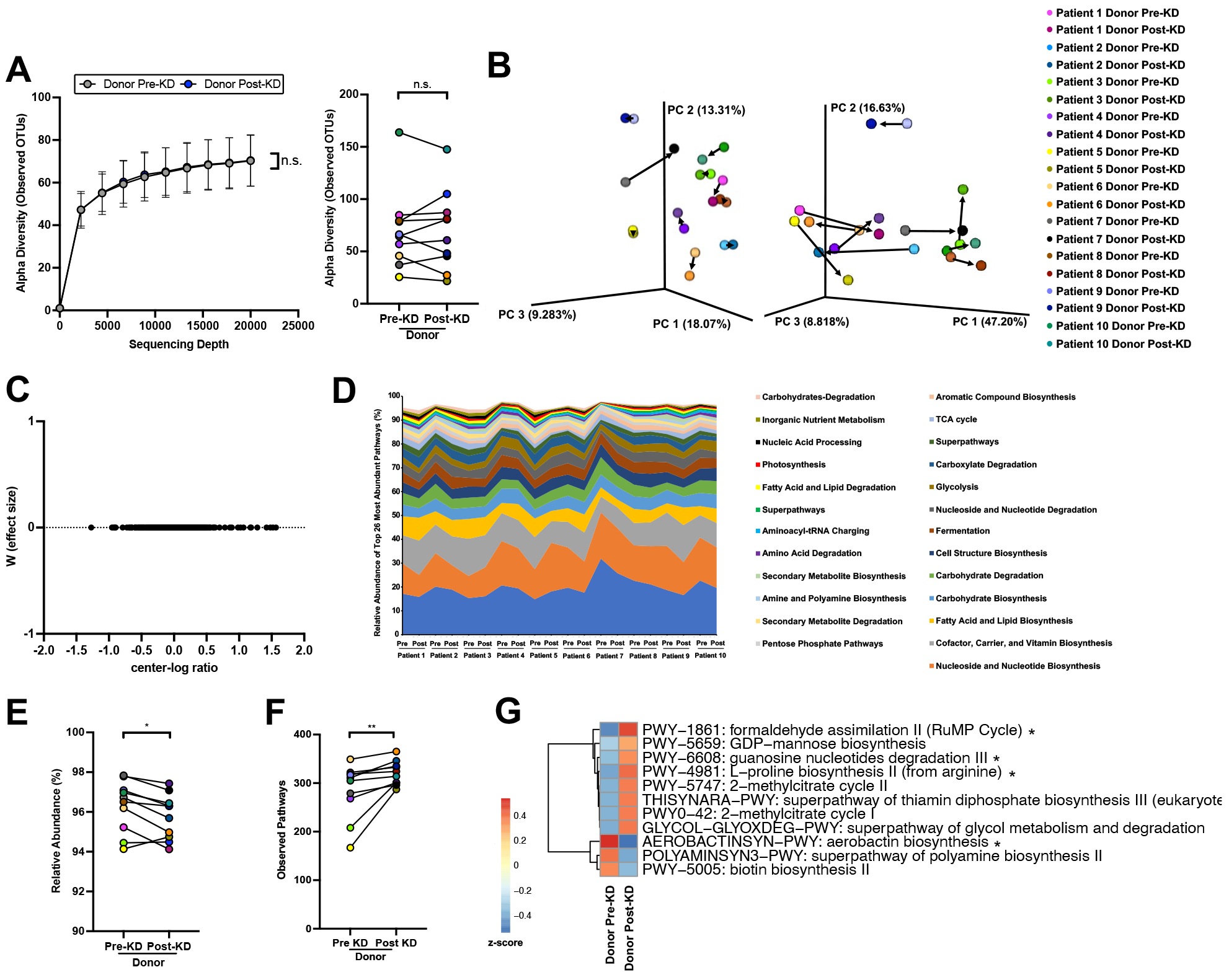
Clinical KD is associated with alterations in the functional potential, but not composition, of the gut microbiota in a cohort of children with refractory epilepsy, Related to Figure 1. **(A)** Alpha diversity as measured by rarefaction curve (left) and observed OTUs (right) of matched donor pre-KD (n=10) and post-KD (n=10) fecal microbiota samples showing no difference in alpha-diversity. (two-way ANOVA with Sidak (left); two-tailed, Wilcoxon matched-pairs signed rank test (right)). **(B)** Principal coordinate analysis of unweighted (left) and weighted (right) UniFrac distances from 16S rRNA gene sequencing of donor pre-KD (n=10) and post-KD (n=10) fecal microbiota samples shifting composition when introduced to the clinical KD. **(C)** ANCOM taxonomic differential abundance testing displaying no differentially abundant taxa by the W score (effect size) metric when comparing donor pre-KD (n=10) and post-KD (n=10). **(D)** Total composition per human donor sample of the top 26 MetaCyc superclass metagenome functional pathways accounting for >94% of relative abundance, for each donor pre-KD (n=10) and post-KD (n=10) fecal microbiota samples. **(E)** Difference in total abundance of the 26 most abundant pathways between matched donor pre-KD (n=10) and post-KD(n=10) fecal microbiota samples (two-tailed, Wilcoxon matched-pairs signed rank test). **(F)** Total number of observed MetaCyc functional pathways in matched donor pre-KD (n=10) and post-KD (n=10) fecal microbiota samples (two-tailed, Wilcoxon matched-pairs signed rank test). **(G)** Heatmap displaying differentially abundant MetaCyc functional pathways associated with donor post-KD (n=10) relative to pre-KD (n=10) by MaAsLin2 analysis with a p-value < 0.1. (pathways with a p-value <0.05 are denoted with *). Data is displayed as mean ± SEM, unless otherwise noted. *p < 0.05, **p < 0.01; KD, ketogenic diet; n.s, no statistical significance.

**Figure S2:**
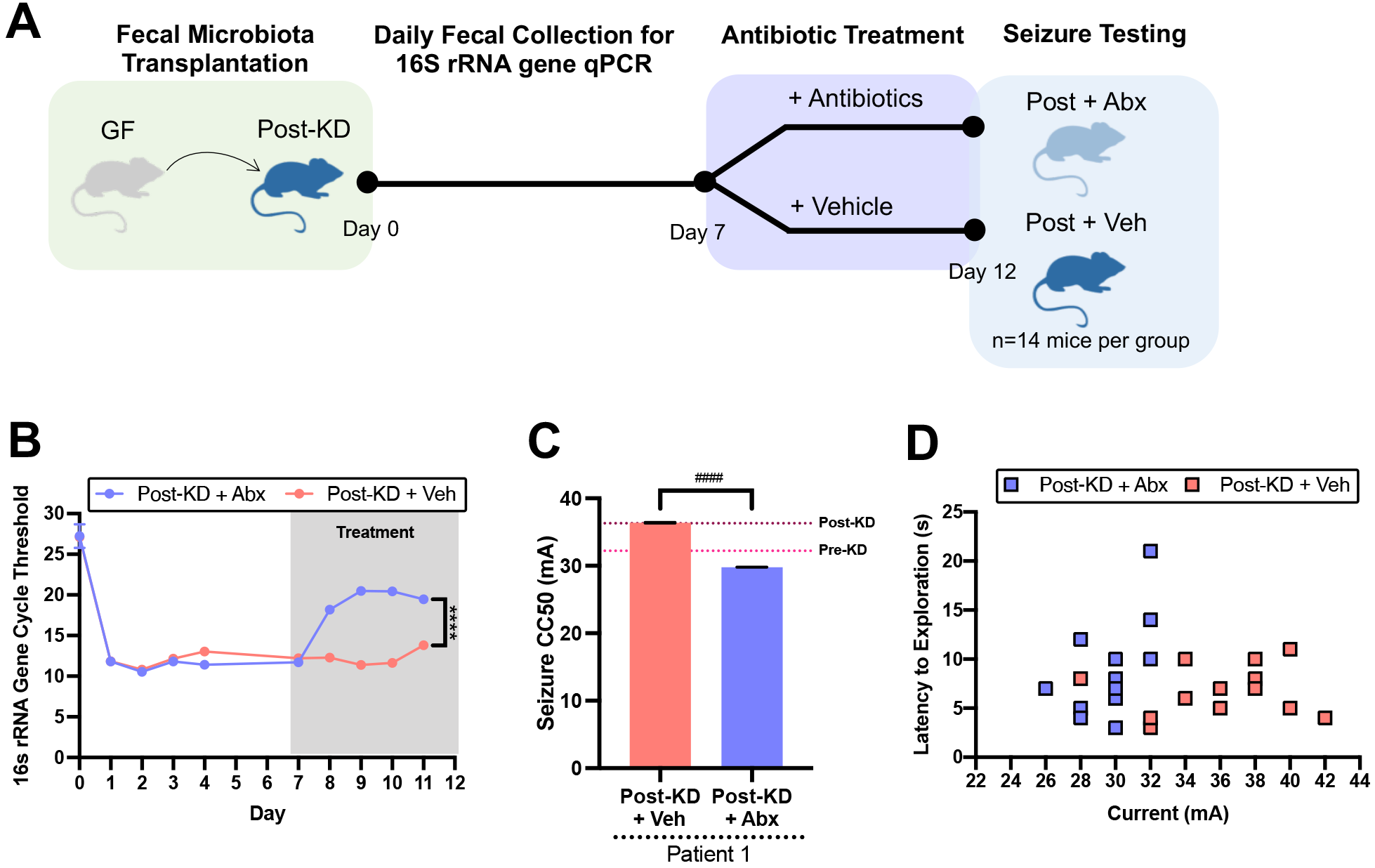
Antibiotic treatment abrogates the seizure protective effects of inoculation with the clinical KD-associated human gut microbiome, Related to Figure 1. **(A)** Experimental schematic for transfer of human donor fecal microbiota samples to germ-free (GF) mice, followed by 5 days of oral antibiotic (Abx) or vehicle (Veh) treatment, and then 6-Hz psychomotor seizure testing. **(B)** Bacterial loads as measured by quantitative PCR of the 16S rRNA gene from fecal pellets collected once daily before and during Abx or Veh treatment (two-way ANOVA with Sidak, n=3 cages of 3 mice each). **(C)** 6-Hz seizure thresholds for mice inoculated with patient 1 post-KD human microbiota treated with Abx (n=12) or Veh (n=14). Reference lines denote seizure thresholds for mice inoculated with patient 1 post-KD and pre-KD relative control fecal microbiota from Figure 1B (One-way ANOVA with Tukey’s, with # denoting statistical differences when considering within-patient recipient mice as technical replicates). **(D)** Latency to exploration for each Abx (n=12) and Veh (n=14) mouse that underwent 6-Hz psychomotor seizure testing. Data is displayed as mean ± SEM, unless otherwise noted. \*\*\*\**p* < 0.0001. ^####^*p* <0.0001 (for within-patient mouse recipients).

**Figure S3:**
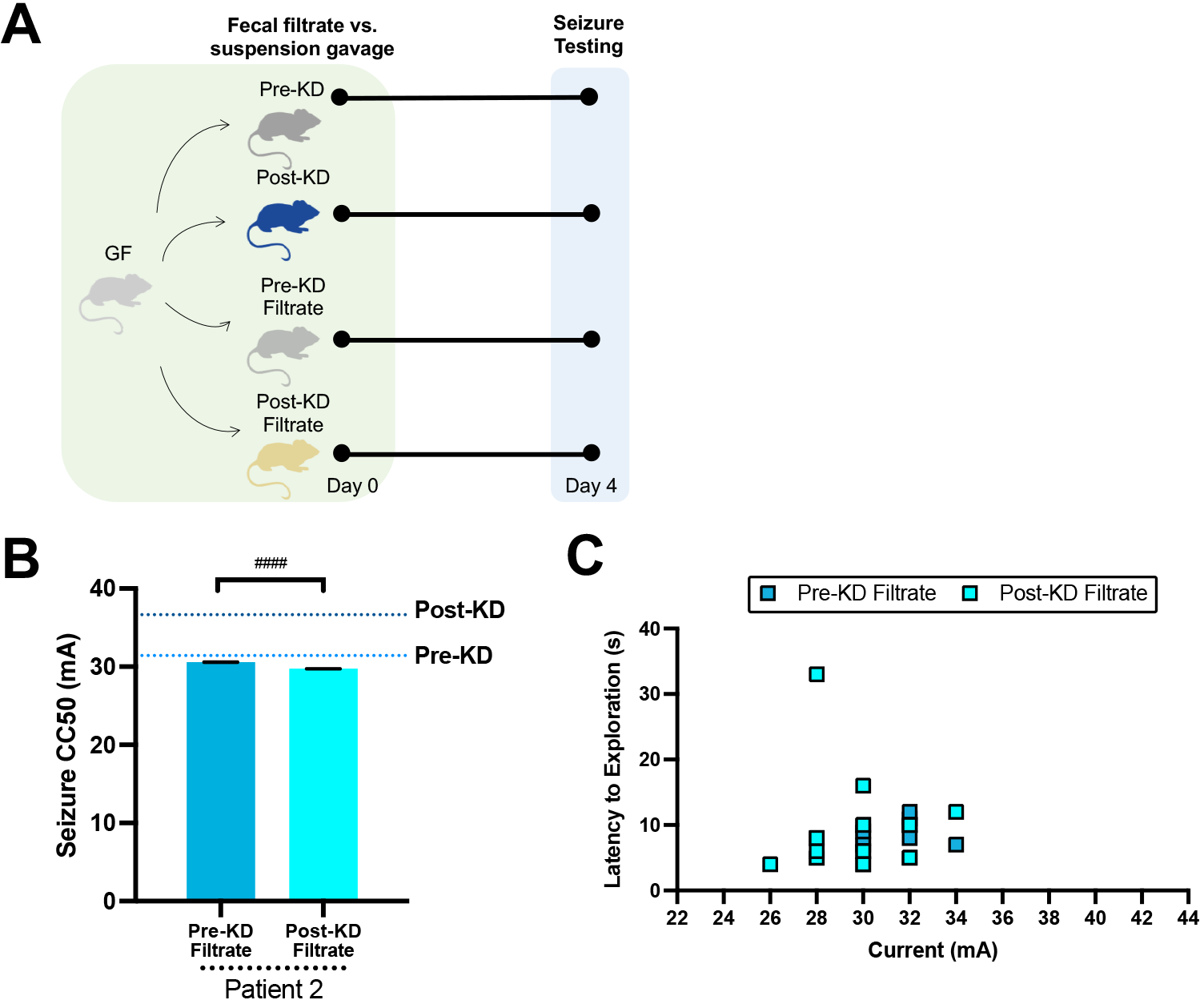
Sterile filtration prevents the seizure protective effects of transfer of the clinical KD-associated human gut microbiome, Related to Figure 1. **(A)** Experimental design for administration of human donor fecal filtrate samples to germ-free (GF) mice, followed by 6-Hz seizure testing 4 days later. **(B)** Seizure thresholds for mice treated with sterile filtered pre-KD (n=14) and sterile filtered post-KD (n=13) fecal samples. Reference lines denote seizure thresholds for mice transplanted with unfiltered patient 2 post-KD and pre-KD relative control fecal microbiota from Figure 1B (One-way ANOVA with Tukey’s, with # denoting statistical differences when considering within-patient recipient mice as technical replicates). **(C)** Latency to exploration for mice treated with sterile filtered pre-KD (n=14) and sterile filtered post-KD (n=13) that underwent 6-Hz psychomotor seizure testing. Data is displayed as mean ± SEM, unless otherwise noted. ^####^ p <0.0001 (for within-patient mouse recipients).

**Figure S4:**
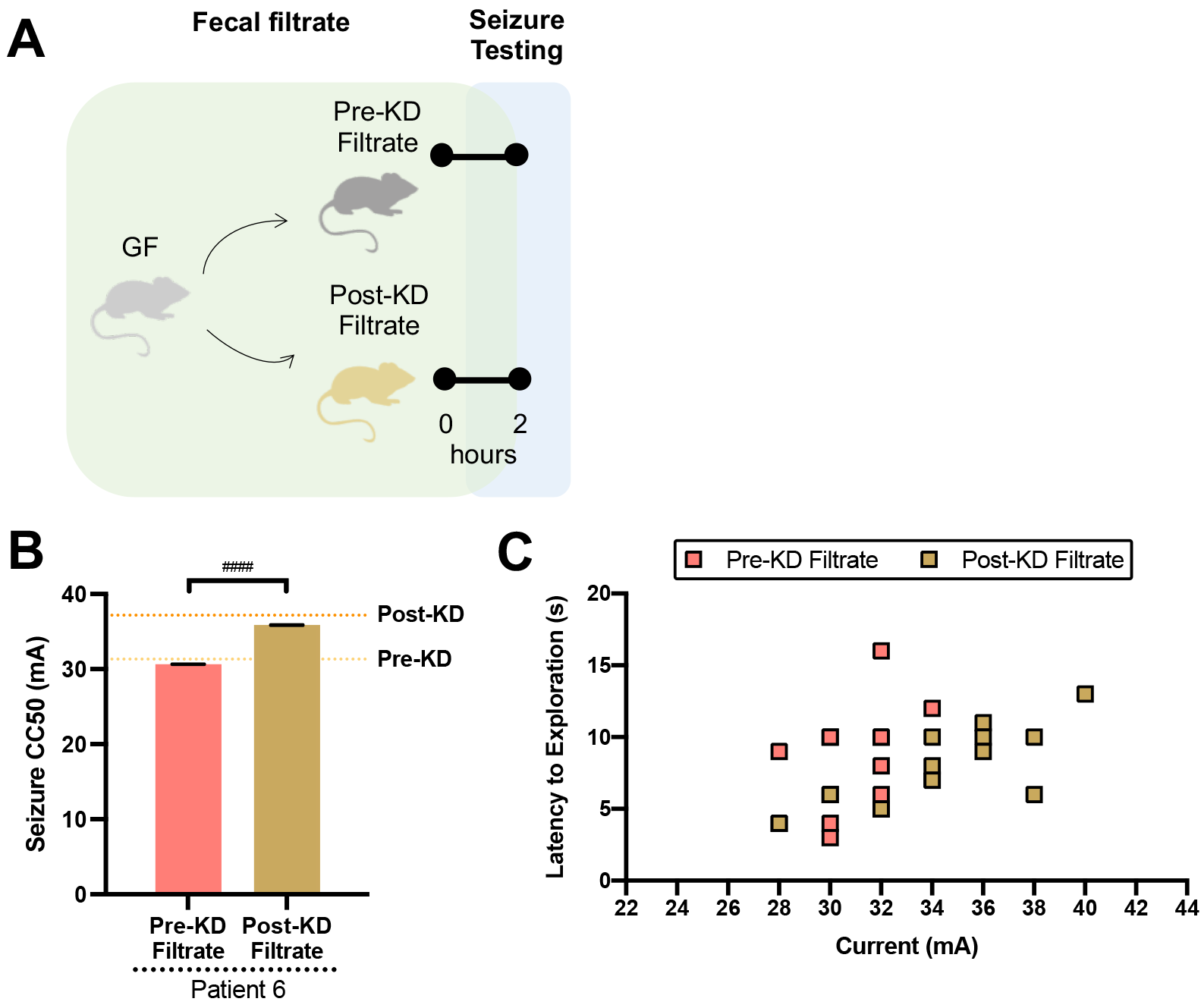
Small molecules from the clinical KD-associated human gut microbiome confer acute seizure protection, Related to Figure 1. **(A)** Experimental design for administration of human donor fecal filtrate samples to germ-free (GF) mice, followed by 6-Hz seizure testing 2 hours later. **(B)** Seizure thresholds for mice treated with sterile filtered pre-KD filtrate (n=13) and sterile filtered post-KD filtrate (n=14) fecal samples. Reference lines denote seizure thresholds for mice transplanted with unfiltered patient 6 post-KD and pre-KD relative control fecal microbiota from Figure 1B (One-way ANOVA with Tukey’s, with # denoting statistical differences when considering within-patient recipient mice as technical replicates). **(C)** Latency to exploration for mice treated with sterile filtered pre-KD (n=13) and sterile filtered post-KD (n=14) that underwent 6-Hz psychomotor seizure testing. Data is displayed as mean ± SEM, unless otherwise noted. ^####^ p <0.0001 (for within-patient mouse recipients).

**Figure S5:**
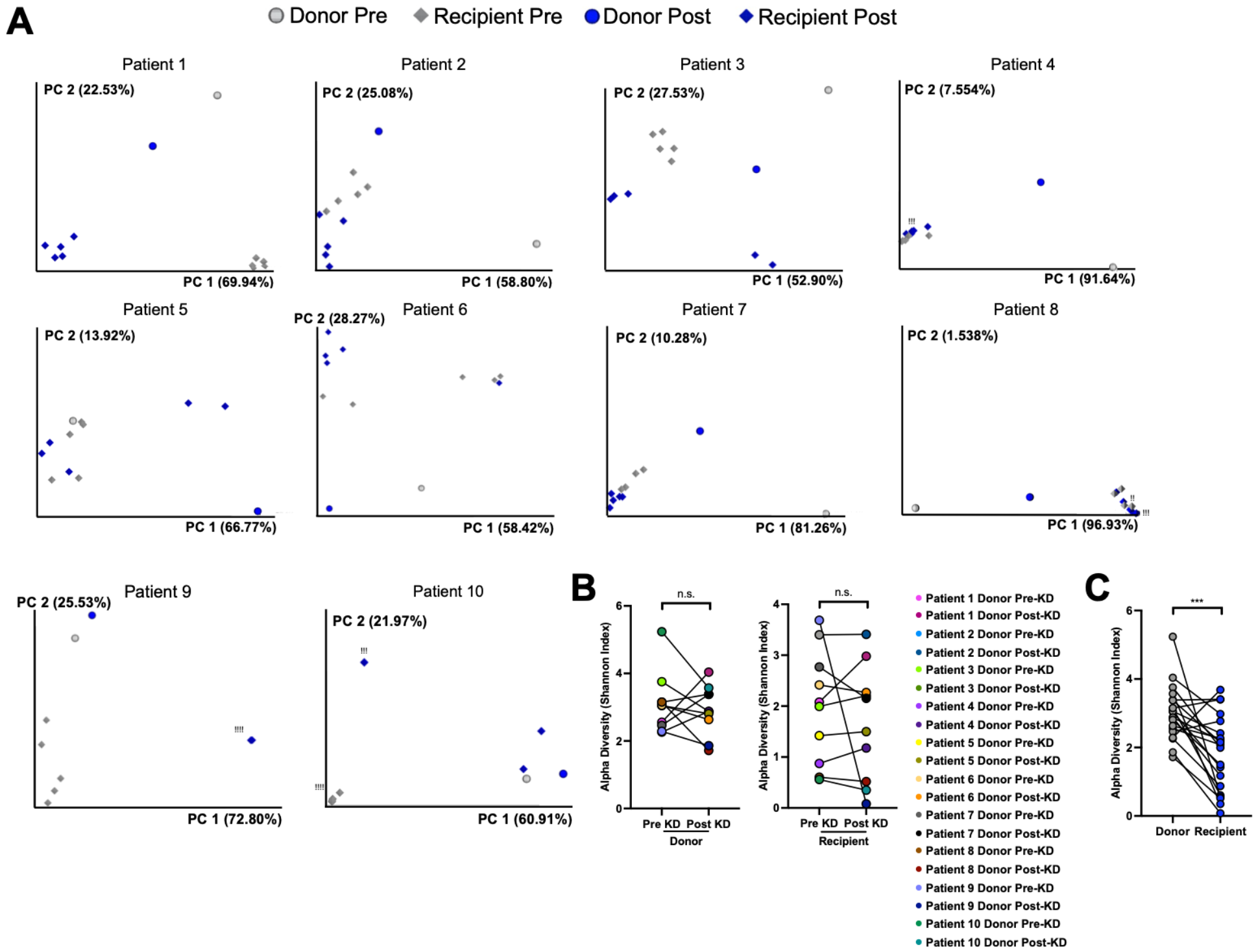
Taxonomic fidelity of human microbiota transfer to mice, Related to Figure 1. **(A)** Principal coordinates analysis of weighted UniFrac distances from 16S rRNA gene sequencing of fecal samples from matched human donors and mouse recipients (for each graph: n = 1 donor patient (10 patients total), 4-5 recipient cages of recipient mice per pre-KD vs. post-KD condition, ! = 1 overlapping data point not visible). **(B)** Shannon index alpha-diversity of fecal microbiota from human donor pre-KD and post-KD samples (left) and matched mouse recipient pre-KD and post-KD samples (right) (two-tailed, Wilcoxon matched-pairs signed rank test, donors: n=10 patients, recipients: n=10 per patient condition, where each n is an average from 4-5 cages per patient). **(C)** Shannon index alpha-diversity of fecal microbiota from all human donor samples (n=20 patients) and all matched mouse recipient samples (two-tailed, Wilcoxon matched-pairs signed rank test; n=20 patient conditions, where each n is an average from 4-5 cages per patient). Data is displayed as mean ± SEM, unless otherwise noted. ***p < 0.001, n.s.=not statistically significant.

**Figure S6:**
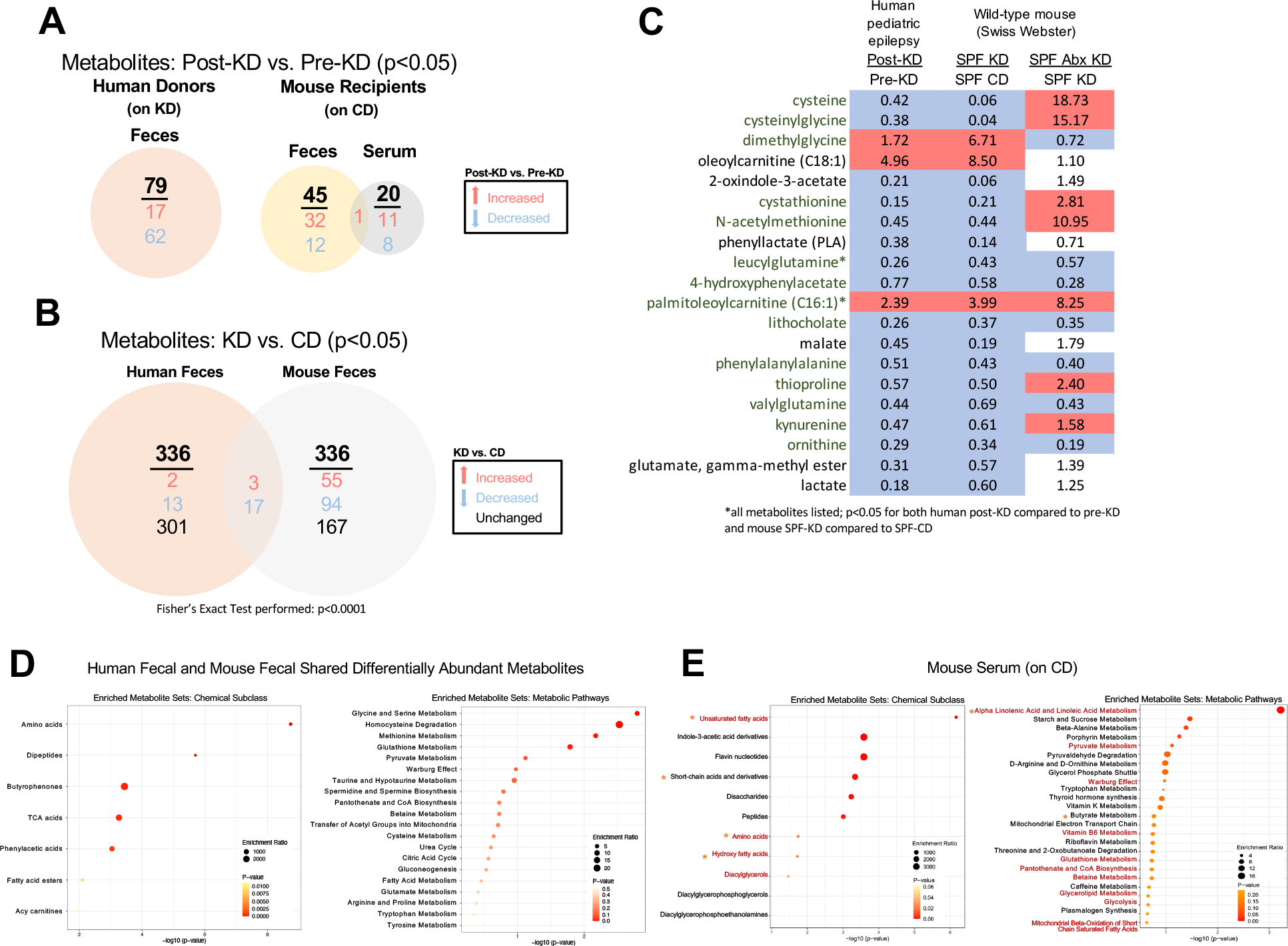
The clinical KD alters metabolomic profiles in human fecal samples and in fecal and serum samples of mice inoculated with human microbiota, Related to Figure 2. **(A)** Differentially abundant metabolites (p<0.05) in post-KD compared to pre-KD samples of human donor feces, mouse recipient feces, and mouse recipient blood (Two-tailed matched pairs Student’s t-test, n=10 per condition, where each recipient sample is pooled from 5 recipient mice per donor patient sample) **(B)** Differentially abundant metabolites (p<0.05) in post-KD compared to pre-KD samples of human donor feces, which were also significantly altered in conventional mice (SPF) fed KD chow or vitamin- and mineral-matched control diet (CD) for 14 days. Red font denotes the subset of metabolites that were further altered by pre-treating KD chow-fed mice with antibiotics (Abx) to deplete gut bacteria. (human: Two-tailed matched pairs Student’s t-test, n=10 per condition; mouse: ANOVA contrasts, n=8 per condition). **(C)** Differentially abundant metabolites (p<0.05) in human feces (post-KD compared to pre-KD) and feces of mice fed KD vs. CD chow for 14 days (Human fecal: Two-tailed matched pairs Student’s t-test, n=10 per condition, where each recipient sample is pooled from 5 recipient mice per donor patient sample; Mouse fecal: two-way ANOVA with contrasts, n=8 per condition; Fisher’s Exact Test). **(D)** Metabolite set enrichment analysis of chemical subclass for the 20 differentially abundant metabolites (p<0.05, matched pairs Student’s t-test) found in both human post-KD vs pre-KD fecal samples and SFP mouse KD vs CD fecal samples (left) (human: n=10 per condition, where each sample is pooled from 5 recipient mice per donor patient sample; mouse: n=8 per condition). Metabolite set enrichment analysis of SMPDB pathways for the 20 differentially abundant metabolites (p<0.05, matched pairs Student’s t-test) found in both human post-KD vs pre-KD fecal samples and SFP mouse KD vs CD fecal samples (right) (human: n=10 per condition, where each sample is pooled from 5 recipient mice per donor patient sample; mouse: n=8 per condition). **(E)** Metabolite set enrichment analysis of chemical subclass for differentially abundant metabolites (p<0.05, matched pairs Student’s t-test) in recipient mouse post-KD vs pre-KD serum samples (left) (n=10 per condition, where each sample is pooled from 5 recipient mice per donor patient sample). Metabolite set enrichment analysis of SMPDB pathways for differentially abundant metabolites (p<0.05, matched pairs Student’s t-test) in recipient mouse post-KD vs pre-KD serum samples (right) (n=10 per condition, where each sample is pooled from 5 recipient mice per donor patient sample). Red font denotes metabolic pathways altered in post-KD vs pre-KD mouse serum that are shared with those differentially regulated in post-KD vs pre-KD mouse feces and/or human feces. Orange asterisks (*) denote additional chemical subclasses that are relevant to KD based on existing literature. KD, ketogenic diet; SPF, specific pathogen free conventionalized mice; CD, control diet; Abx, antibiotic SMPDB, The Small Molecule Pathway Database

**Figure S7:**
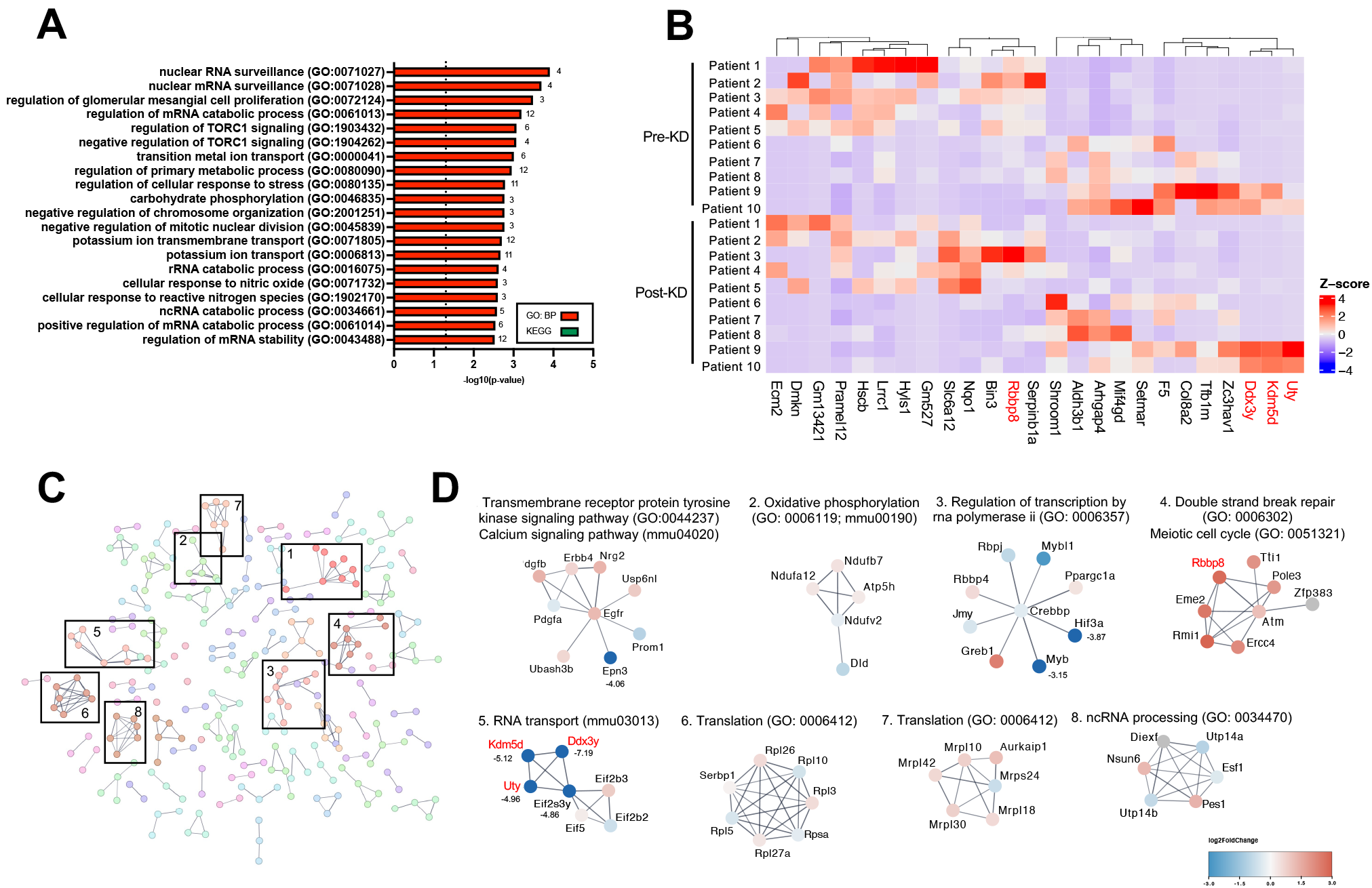
Mice inoculated with the post-KD microbiota exhibit alterations in the frontal cortical transcriptome, Related to Figure 3. **(A)** GO: Biological Process gene ontology of differentially expressed genes (p<0.05) in recipient mouse post-KD compared to pre-KD frontal cortex samples, top 20 ranked by p-value (n=10 per patient diet condition, where each sample is pooled from 6 recipient mice per donor patient sample). **(B)** Heatmap of top 25 differentially expressed genes in recipient mouse post-KD compared to pre-KD frontal cortex ranked by p-value, smallest to largest, with log2-fold change >2 (n=10 per patient diet condition, where each sample is pooled from 6 recipient mice per donor patient sample). **(C)** Protein interaction network with MCL clustering based upon mouse recipient post-KD and pre-KD frontal cortex transcriptomics which appeared in both GO and STRING network enrichment analyses, STRING network enrichment score >0.7 (n=10 per patient diet condition, where each sample is pooled from 6 recipient mice per donor patient sample). **(D)** Functional enrichment of top MCL sub-network clusters from frontal cortex transcriptomics STRING network analysis, proteins are colored based on their overall log2FC. If log2FC >3 or <-3, the value is listed next to the node name (n=10 per patient diet condition, where each sample is pooled from 6 recipient mice per donor patient sample). KD, ketogenic diet; GO, gene ontology; MCL, Markov Cluster Algorithm.

